# Contrastive Learning-Enhanced Drug Metabolite Prediction Drives the Discovery of an Orally Bioavailable RSK4 Inhibitor for Esophageal Cancer

**DOI:** 10.1101/2025.06.02.657523

**Authors:** Huan He, Manzhan Zhang, Xiaoxiao Yang, Shuai He, Xiayu Shi, Feng Hu, Chang Liu, Xingsen Zhang, Na Chen, Xiaoqian Zhu, Leihao Zhang, Tianyu Ye, Rong Zhang, Yanru Yang, Rui Wang, Zhenjiang Zhao, Zhuo Chen, Xuhong Qian, Honglin Li, Zhe Wang, Kai Zhang, Kangdong Liu, Shiliang Li

**Affiliations:** School of Pharmacy, East China University of Science & Technology, Shanghai 200237, China; Innovation Center for AI and Drug Discovery, School of Pharmacy, East China Normal University, Shanghai 200062, China; Department of Pathophysiology, School of Basic Medical Sciences, Zhengzhou University, Zhengzhou, Henan, 450001, China; State Key Laboratory of Holistic Integrative Management of Gastrointestinal Cancers, Department of Pathology, Xijing Hospital and School of Basic Medicine, Fourth Military Medical University, Xi’an, 710032, China; Department of Pathology, School of Basic Medicine and Forensic Science, Baotou Medical College, Baotou, 014060, China

**Author notes:** Authors to whom correspondence should be addressed. Zhe Wang, Kai Zhang, Kangdong Liu, Shiliang Li. These authors contributed equally (Huan He, Manzhan Zhang, Xiaoxiao Yang, Shuai He).

## Abstract

Drug metabolism liabilities are a major bottleneck in drug development, necessitating accurate metabolite prediction tools. While AI-driven methods have advanced, challenges remain in achieving high-precision, robustness, broad reaction coverage and seamless integration into the drug development workflow. To address this, we first constructed the most comprehensive human-specific drug metabolism database to date, encompassing 11,665 human metabolic reactions and 2,497 high-quality rules. Leveraging this resource, we developed ConMeter (**Con**trastive learning and feature interaction based **Met**abolic **R**eaction prediction), a novel model integrating chemical feature interaction and contrastive learning. ConMeter demonstrates significant improvements, achieving ∼80% accuracy in identifying at least one metabolite within the top 5 predictions and showing an average 25% relative enhancement in ranking true positives compared to state-of-the-art methods. The comprehensive database also expands the range of predictable metabolic reactions. To demonstrate the utility of ConMeter, we integrated it into a metabolite prediction-guided drug design strategy aimed at addressing the low oral bioavailability of a metabolically vulnerable RSK4 lead compound — a pervasive issue among known RSK inhibitors. This strategy successfully yielded R636, a novel RSK4 inhibitor exhibiting remarkably improved bioavailability (a 63-fold increase, reaching 63% absolute bioavailability) while maintaining its potent activity. Notably, R636 showed favorable safety profile and significant anti-tumor efficacy against esophageal squamous cell carcinoma (ESCC) *in vitro* and in two distinct patient-derived xenograft (PDX) mouse models. This work introduces ConMeter as a powerful and practical tool for high-accuracy metabolite prediction, demonstrating its potential to accelerate lead optimization, exemplified by the discovery of R636 as a promising orally bioavailable candidate for ESCC.

## INTRODUCTION

Drug discovery is a lengthy and costly process, often taking up to a decade with clinical trials lasting 6 to 8 years and costs exceeding $1 billion^1–3^. A major contributor to the high attrition rates and escalating costs in drug discovery is the emergence of unexpected pharmacokinetic (PK) issues, particularly those arising from drug metabolism, which often remain undetected until late-stage development^2, 4–7^. Drug metabolism, a critical component of a drug’s ADMET (Absorption, Distribution, Metabolism, Excretion, and Toxicity) profile, plays a central role in determining its PK characteristics by mediating activation, inactivation, detoxification, and potential toxicity^8^. For orally administered drugs, effectively overcoming first-pass metabolism in the liver is especially crucial for achieving adequate systemic exposure and bioavailability^9–11^. Consequently, the accurate prediction and comprehensive understanding of a drug candidate’s metabolic fate are not only advantageous but essential for optimizing its PK properties and ensuring successful clinical translation^12–16^.

In the optimization of new drugs, a common strategy employed by researchers is the modification of the site of metabolism (SOM) in lead compounds to enhance overall PK properties, particularly oral bioavailability and half-life^12–16^. While preclinical studies, typically conducted in animal models, offer valuable preliminary insights into drug candidate metabolism, a critical limitation arises from the significant interspecies variations in enzyme expression, activity, and substrate specificity^17^. These inherent differences between animal and human metabolic profiles present a major impediment to accurately predicting human pharmacokinetics and drug efficacy based on preclinical data. Furthermore, the study of metabolites in humans during clinical research is not only resource-intensive but also demands considerable specialized expertise^12, 13^. These combined factors underscore the intrinsic complexity of developing drug candidates with optimal metabolic profiles, thereby highlighting the urgent need for innovative tools and methodologies to accelerate drug discovery and development. To address this need, *in silico* metabolite prediction methods, such as SyGMa^18^, BioTransformer^19^, and GLORYx^20^, have emerged. However, these early methods often suffer from limitations in scalability and generalizability. More recently, MetaTrans^21^, an AI-driven approach using SMILES for sequence translation, demonstrated improved metabolite detection, particularly for reactions involving rare enzymes. Despite these advancements, existing tools still struggle to effectively discriminate between metabolized and non-metabolized compounds, resulting in high false positive rates and limited generalizability. Moreover, a key shortcoming of current approaches lies in their failure to fully capture the intricate relationships within metabolic reactions. They often treat reactants and products as discrete chemical entities, neglecting the crucial structural correlations between them. This fundamental limitation underscores the continued necessity for developing more sophisticated algorithms that can significantly improve the predictive accuracy of metabolic properties and drive further progress in this critical research domain.

To overcome the aforementioned limitations in current metabolite prediction approaches, we introduce ConMeter, a novel and robust metabolite prediction model. ConMeter is built upon a comprehensive metabolic reaction dataset and utilizes an innovative template extraction algorithm. This algorithm is crucial for generating high-quality positive and negative sample pairs, which are essential for effective model training. Furthermore, ConMeter employs advanced feature extraction techniques meticulously designed to capture the intrinsic structural correlations between reactants and their corresponding metabolites. By harnessing the power of contrastive learning, ConMeter is specifically trained to effectively differentiate between valid and invalid reactant-metabolite pairs. This unique combination of chemical features and learning strategy ensures improved model performance, enhanced scalability, and broader generalizability. Notably, the model achieved remarkable accuracy at TOP 5 (78.38% and 87.50%) and TOP 10 (86.49% and 96.88%) to identify at least one metabolite in two distinct test sets. ConMeter exhibited superior performance across all key metrics (TOP5, 10, 13, 20), demonstrating significant improvements in true positive metabolite identification with average gains of 27% and 24% over existing tools in the two aforementioned test sets, respectively.

Esophageal squamous cell carcinoma (ESCC) ranks among the most lethal malignancies worldwide, characterized by rapid progression and poor prognosis. Despite therapeutic advancements, persistent challenges, including late-stage diagnosis, reliance on complex multimodal therapies, and limited targeted treatments, contribute to dismal 5-year survival rates, underscoring the urgent need for novel therapeutic strategies^22^. Ribosomal S6 kinase 4 (RSK4), a serine/threonine kinase in the ribosomal S6 kinase family^23–27^, is now recognized as a critical driver of cancer stem cell-like (CSC) properties and a mediator of radiation resistance in ESCC^28^. Compelling evidence indicates that the RSK4 inhibitor BI-D1870 can significantly reduce CSC traits and enhance radiosensitivity in preclinical ESCC models, solidifying RSK4 as a highly promising therapeutic target for ESCC^29^. In our prior work, we identified a potent small-molecule RSK4 inhibitor, compound 14f, which exhibited exceptional anti-ESCC activity both *in vitro* and *in vivo*^30^. However, mirroring a significant limitation observed across known RSK inhibitors such as BI-D1870^31^, LJH685^32^ and LJI308^33^ (**Extended Data Fig. 1**), 14f suffered from profoundly inadequate oral bioavailability (*F* = 0.99%). A critical structural feature shared by these compounds, including 14f, is a phenolic hydroxyl moiety, which serves as a key pharmacophore essential for forming crucial intermolecular interactions with the RSK4 receptor, thereby driving potent inhibitory activity at the molecular level. Paradoxically, this essential phenolic hydroxyl group, while crucial for efficacy, is highly susceptible to rapid hepatic phase II conjugation, leading to accelerated elimination and significantly limiting oral bioavailability due to substantial first-pass metabolism^34, 35^. Therefore, the successful development of an orally bioavailable RSK4 inhibitor for ESCC therapy critically hinges on overcoming the bioavailability challenges inherent to this class of compounds.

In this study, we strategically employed metabolite prediction guided drug design as a rational approach to optimize the metabolic properties of 14f. This approach culminated in the successful identification of R636, a novel drug candidate exhibiting significantly improved oral bioavailability (*F* = 63%) without compromising target potency. Subsequent *in vitro* and *in vivo* evaluations of R636 unequivocally demonstrated its robust anti-ESCC efficacy, especially in PDX mouse models. In conclusion, this work not only delivers ConMeter as a valuable and innovative tool for advancing metabolite prediction, but also, perhaps more importantly, identifies R636 as a highly promising orally bioavailable therapeutic candidate poised for further development in the fight against ESCC.

## RESULTS

### A comprehensive, human-specific drug metabolism database with broad reaction rules coverage

Our predicting model is designed on the fundamental principle that drug metabolism involves *in vivo* biotransformation, resulting in metabolites with distinct structural identities from their parent drugs (**Fig. 1a**). To this end, we elaborately compiled a dataset encompassing 11,665 human metabolic reactions sourced from a variety of reputable database for training the AI model (**Fig. 1b**). This extensive collection, categorized into 17 distinct reactions types, represents the most comprehensive human-specific drug metabolism database to date, characterized by its extensive coverage of diverse reaction rules (**Fig. 1c**).

**Fig. 1.**
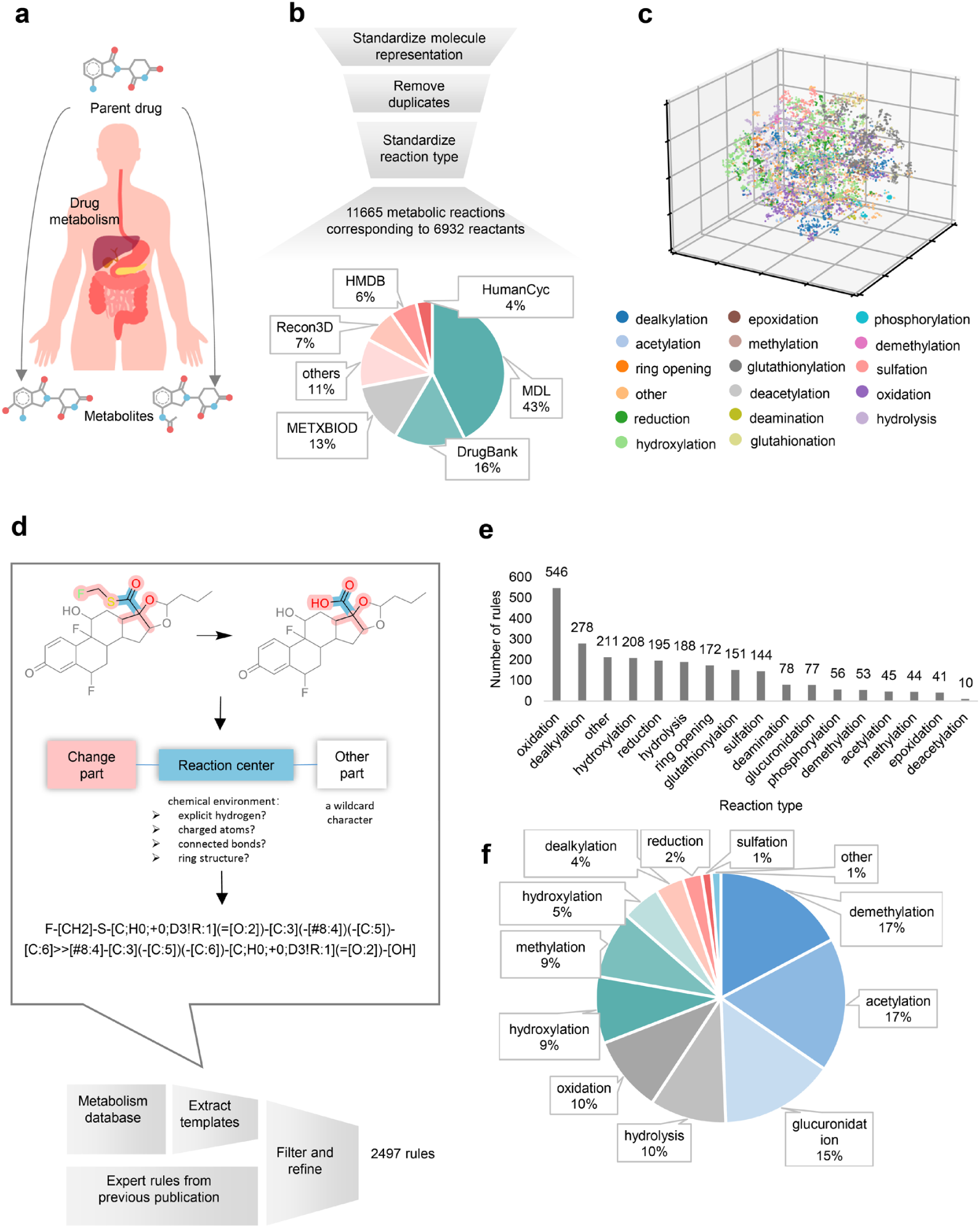
A comprehensive, human-specific drug metabolism database with broad reaction rules coverage. **a**, Schematic diagram of drug metabolism in the human body. **b**, Integration of multi-source data for the construction of a comprehensive human metabolism database. The sources of the database are analyzed by pie chart statistics. **c**, 3D t-SNE visualization of the 11665 metabolic reactions based on molecular fingerprint, different colors represent different types of reactions. **d**, Development and refinement of the templates database via a novel template extraction strategy based on a forward enumeration algorithm. **e**, A statistical diagram of the reaction types of the template database. **f**, Statistics of metabolic reaction types in our newly curated test set 2.

Recognizing the inherent limitations of conventional rule-based methods in terms of scalability and generalizability, we developed a novel template extraction algorithm leveraging the capabilities of forward enumeration algorithm^36^ and RDKit^37^(**Fig. 1d**). Applying this algorithm, we initially extracted a substantial number of reaction rules, inevitably containing redundancy. Subsequently, we implemented a rigorous filtering and manual refinement process, ultimately yielding a set of 2497 high-quality, non-redundant rules. This refined reaction rule set ensures significantly enhanced coverage of metabolic transformations (**Fig. 1e**).

To rigorously evaluate the predictive performance of ConMeter, we assembled two independent test sets. Building upon the GLORYx test set (test set 1), which includes 136 reactions from 37 reactants, we further curated a novel and contemporary test set (test set 2). This set was meticulously compiled from metabolic reactions documented in the recent literature (2019-2023), encompassing 104 reactions from 31 reactants. Detailed information and references for test set 2 are provided in **Supplementary Data 1**, and the distribution of reaction types within test set 2 is depicted in **Fig. 1f**.

### Metabolite prediction model ConMeter

To effectively distinguish true metabolites from false positives, we developed ConMeter, a deep learning framework that synergistically integrates chemical feature interaction and contrastive learning algorithms (**Fig. 2**). ConMeter leverages a template-based approach to generate metabolite candidates (**Fig. 2a**). By precisely matching reactants with appropriate reaction templates, we can generate authentic metabolite structures, thereby getting high-confident positive pairs for model training. Furthermore, to ensure robust model learning and discrimination, we devised a strategy to construct a substantial dataset of negative pairs. This is achieved by systematically pairing reactants with non-corresponding reaction templates, while meticulously maintaining structural feasibility within the generated negative metabolite structures. By enriching both the diversity and quantity of training data, ConMeter is empowered to learn from a wide spectrum of metabolic transformations. The model processed chemical molecular fingerprints of both reactants and products, enabling it to discern the nuanced chemical and structural relationships that fundamentally govern metabolic reactions.

**Fig. 2.**
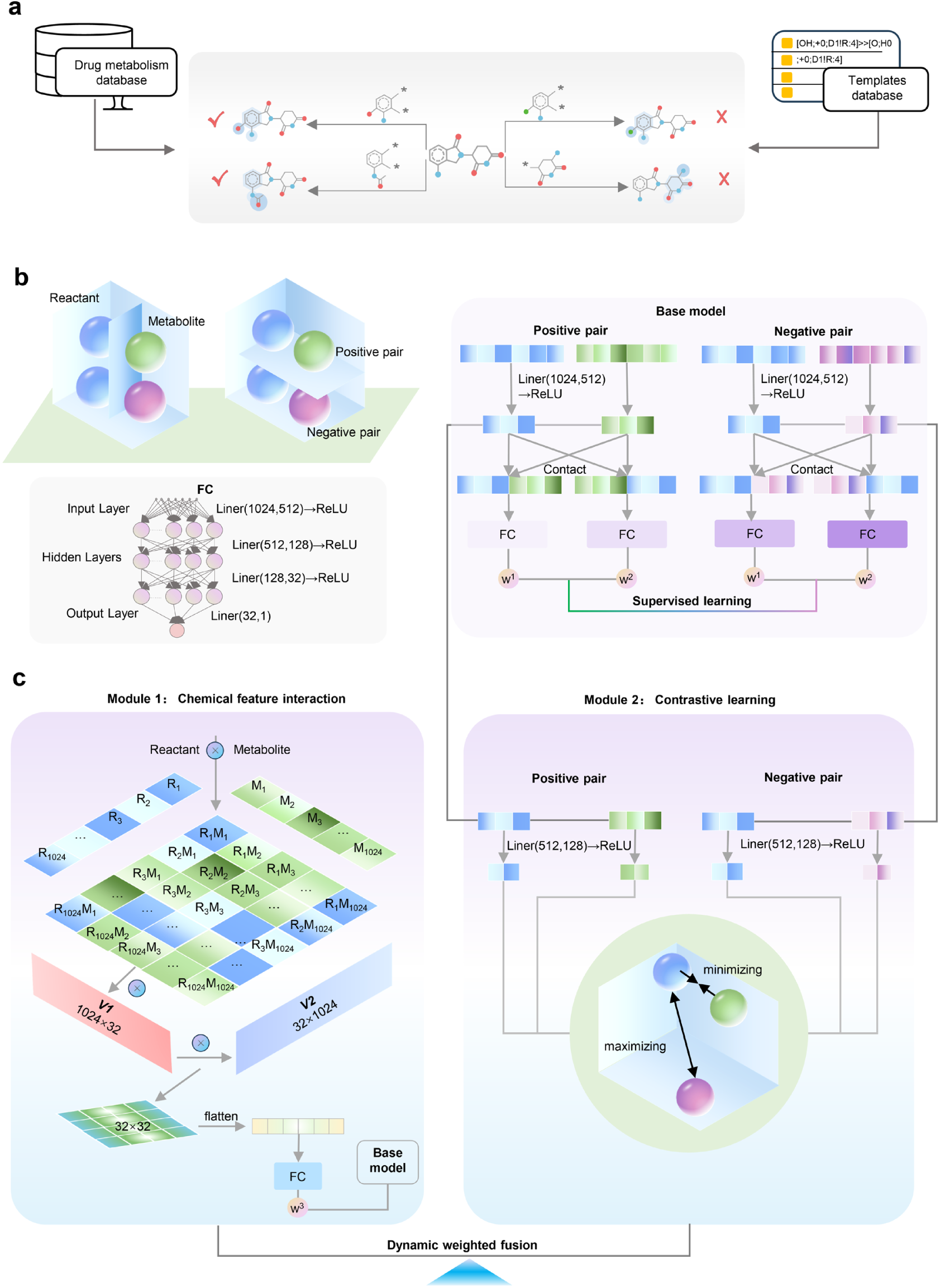
An illustrative diagram of the metabolite prediction model ConMeter. **a**, The matching process for generating positive pairs and negative pairs. **b,** The schematic diagram of the relationship between reactants and metabolites, positive and negative pairs (upper section), and the fundamental framework of ConMeter’s base model (lower section). **c**, The diagram of the metabolite prediction model integrating the chemical feature interaction algorithm and the contrastive learning algorithm. FC: Full connect neural network.

The architecture of the ConMeter framework is schematically depicted in **Fig. 2b, 2c** and comprises distinct yet interconnected modules. Firstly, ConMeter incorporates a Chemical Feature Interaction Module. This module ingeniously employs the outer product of the reactant’s and metabolite’s feature vectors to effectively capture and encode their intrinsic physicochemical interactions. Following a series of matrix transformations, this module outputs a concise one-dimensional vector representation of the interaction space. The Chemical Feature Interaction Module significantly contributes to true positive identification across both test sets, with substantial improvements particularly in TOP1-20 (**Extended Data Fig. 2a,b**). In Test Set 1, this module achieves an average TP improvement of 10.6% over the base model (**Extended Data Fig. 2c**). Similarly, in Test Set 2, an 8.5% average improvement is observed (**Extended Data Fig. 2d**). The precision-recall curves further indicate that feature interaction particularly enhances precision at lower TOP N ranges while consistently improving recall across all TOP N values (**Extended Data Fig. 2e,f**). Secondly, a Contrastive Learning Module is integrated. Within this module, molecular fingerprint vectors of reactants and metabolites are independently projected into low-dimensional latent spaces via dedicated fully connected networks. Subsequently, contrastive learning is applied in this latent space to enforce proximity between positive pairs while maximizing the separation between positive and negative pairs, driven by a carefully formulated contrastive loss function. The latent representations derived from reactants and metabolites are then concatenated and further processed through a fully connected layer to yield another one-dimensional vector. The Contrastive Learning Module delivers remarkable performance gains, particularly at lower TOP N values (**Extended Data Fig. 3a,b**). In Test Set 1, this module achieves an average 8.9% improvement in true positives (**Extended Data Fig. 3c**). In Test Set 2, the average improvement is more modest at 2.7% (**Extended Data Fig. 3d**). The precision-recall analysis shows that contrastive learning maintains higher precision than the base model at critical lower TOP N values while simultaneously improving recall (**Extended Data Fig. 3e,f**). Crucially, the predictive framework employs a novel dual-weighting mechanism to optimally integrate the output vectors from both the Chemical Feature Interaction Module and the Contrastive Learning Module. In particular, the coefficients sum to one. Backpropagation is used to optimize the weighting coefficients and all other model parameters. Through this synergistic integration of chemical feature interaction and contrastive learning, ConMeter was trained to learn highly discriminative features, ensuring robust and reliable differentiation even in the face of complex and inherently ambiguous metabolic scenarios. The merged model consistently outperforms individual modules and the base model across all metrics. The true positive curves (**Extended Data Fig. 4a,b**) show that the merged model achieves the highest values across most TOP N ranges in both test sets. Precision analysis (**Extended Data Fig. 4c,d**) reveals that the merged model maintains an optimal balance between the strengths of both modules. The recall curves (**Extended Data Fig. 4e,f**) demonstrate the merged model’s superiority in identifying relevant metabolites. Most impressively, the heatmaps (**Extended Data Fig. 4g,h**) quantify these improvements, showing that the merged model achieves the highest average improvement over the base model—17.7% for Test Set 1 and 16.9% for Test Set 2—substantially outperforming both the Feature Interaction (14.5% and 15.0%) and Contrastive Learning (13.5% and 5.3%) modules when used independently.

The entire framework undergoes rigorous training and validation via a five-fold cross-validation protocol, with model performance optimized through a joint objective function combining cross-entropy and contrastive loss terms. During the prediction phase, the trained model outputs a probabilistic score for each reactant-metabolite pair, providing a quantitative measure of reaction likelihood. Furthermore, this probabilistic output enables the deduction of the precise SOM for a given query molecule.

### ConMeter achieves superior performance in metabolite prediction compared with existing tools

To rigorously assess the efficacy of ConMeter, we conducted a comprehensive benchmarking study, systematically comparing its performance against four established drug metabolite prediction tools: GLORYx, BioTransformer, MetaTrans, and SyGMa. Our evaluation focused on the prediction of metabolites generated through single-step reactions, employing both test set 1 and test set 2. Across all key performance metrics (TOP 5, 10, 13 and 20), ConMeter consistently outperformed comparative methods in true positive metabolite identification (**Fig. 3a, 3b**), with an average improvement of 27% and 24% over existing tools in test sets 1 and 2, respectively. Notably, at TOP 5 predictions, ConMeter identified 54 and 53 true positives in test sets 1 and 2, respectively, representing substantial gains over GLORYx (15 and 17 more metabolites), SyGMa (6 and 10 more metabolites), and MetaTrans (15 and 18 more metabolites). This metric is particularly relevant in practical drug discovery settings, where identifying even a single major metabolite within a manageable top-ranked list can be highly valuable. Furthermore, analysis of the total number of true positive predictions unequivocally demonstrates that ConMeter’s innovative template matching methodology achieves a demonstrably broader coverage of metabolic reaction types, surpassing existing tools in both test sets.

**Fig. 3.**
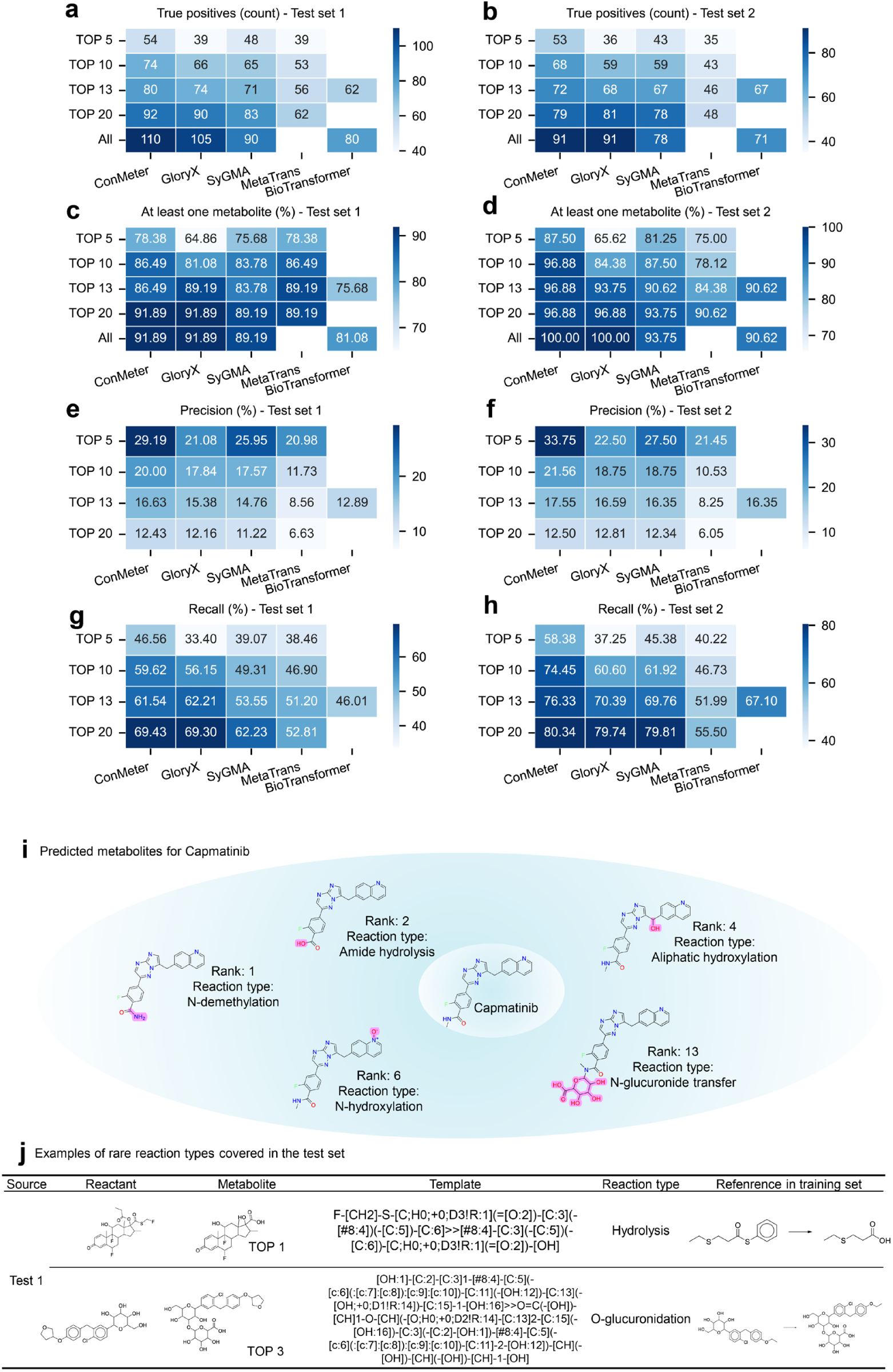
ConMeter achieves superior performance in metabolite prediction compared with existing tools. **a**-**b**, Heatmaps of the number of true positive compounds that are successfully predicted for each reactant in test set 1 (**a**) and test set 2 (**b**) on TOP5, TOP10, TOP13, TOP20 and all predictions. **c**-**d**, Heatmaps of the accuracy that at least one true metabolite is successfully predicted for each reactant in test set 1(**c**) and test set 2 (**d**) on TOP5, TOP10, TOP13, TOP20 and all predictions. **e**-**f**, Heatmaps of precision (**e**) and recall (**f**) for test set 1 compared with the baseline models. **g**-**h**, Heatmaps of precision (**g**) and recall (**h**) for test set 2 compared with the baseline models. **i**, Two examples of prediction results in test set 2 that cover the common reactions. **j**, Examples of rare reaction types covered in the test sets, also displayed the corresponding template extracted by ConMeter.

To gain a more nuanced understanding of ConMeter’s predictive capabilities, we evaluated its accuracy in scenarios where at least one true metabolite is correctly identified within the top-ranked predictions. Specifically, ConMeter demonstrated remarkable accuracy of 78.38% (test set 1) and 87.50% (test set 2) at TOP 5, with an average of 5 (test set 1) and 14 (test set 2) percentage points higher than existing methods. For TOP 10 predictions, ConMeter reached 86.49% and 96.88% accuracy across both test sets, providing strong evidence for its superior predictive capability in practical metabolite identification scenarios (**Fig. 3c, 3d**).

These compelling results not only demonstrated the superior predictive power of the ConMeter model, but also provided valuable guidance for practical application: comprehensive consideration of metabolic sites corresponding to the top 5 or even top 10 predicted metabolites is demonstrably a highly effective strategy for maximizing metabolite identification success. In a direct comparison of recall and precision, ConMeter consistently exhibited superior performance. For precision metrics at TOP 5, ConMeter achieved 29.19% and 33.75% across both test sets, representing absolute average improvements of 6.52 percentage points (test set 1), and 9.93 percentage points (test set 2) over existing methods (**Fig. 3e, 3f**). The recall performance similarly demonstrated ConMeter’s advantages, with TOP 5 values of 46.56% and 58.38%, showing absolute average improvements of 9.58 percentage points (test set 1), and 17.43 percentage points (test set 2) (**Fig. 3g, 3h**). These quantitative improvements clearly demonstrate ConMeter’s significant advancement in mitigating the persistent challenge of high false-positive rates that plagues existing metabolite prediction tools

One of particular note is ConMeter’s remarkable ability to accurately predict both common and less frequent metabolic reactions. For instance, the N-demethylation, Amide hydrolysis, Aliphatic hydroxylation, N-hydroxylation, N-glucuronide transfer of Capmatinib^38^ were all correctly predicted within the Top 13 (**Fig. 3i**). Beyond these common reactions, ConMeter also demonstrated a noteworthy capacity to capture rarer, yet pharmacologically significant, reaction types. This is exemplified by its successful prediction of the hydrolysis of fluticasone propionate (FTP) to fluticasone 17β-carboxylic acid^39^ (**Fig. 3j**), a transformation often challenging for traditional rule-based methods. Furthermore, ConMeter’s capabilities extend to even more specialized reactions, such as glucuronidation, showcasing its versatility across a broad spectrum of metabolic transformations.

In summary, our comprehensive evaluation unequivocally establishes ConMeter’s superior performance, not only in terms of metabolite prediction accuracy but also in its significantly enhanced coverage of diverse metabolic reaction types, surpassing the capabilities of existing state-of-the-art tools.

### ConMeter is a key driver in the discovery of novel, potent and orally bioavailable RSK4 inhibitor R636

Our research group has previously identified that RSK4 is a promising therapeutic target for the treatment of ESCC^28, 30^. Building upon this foundation, and employing structure-based drug design, we successfully discovered lead compound 14f, which exhibited remarkable anti-ESCC activity in both *in vitro* and *in vivo* models. However, despite its potent efficacy, 14f was unfortunately hampered by a critical limitation: an unacceptably high clearance rate and extremely poor oral bioavailability (*F* = 0.99%)^30^, thus impeding its further clinical translation. Based on our prior experience and a preliminary assessment, we hypothesized that rapid metabolism in mice was a primary contributor to the poor bioavailability of 14f, with the phenolic hydroxyl moiety and carbamate moiety identified as potential metabolic hotspots. Subsequently, in an attempt to circumvent these metabolic liabilities, we employed traditional drug modification strategies, leading to the synthesis of compound R632 through scaffold hopping (**Fig. 4a**). Regrettably, this conventional approach proved insufficient, as R632 exhibited only marginally improved, and still critically inadequate, oral bioavailability (*F* =1.2%).

**Fig. 4.**
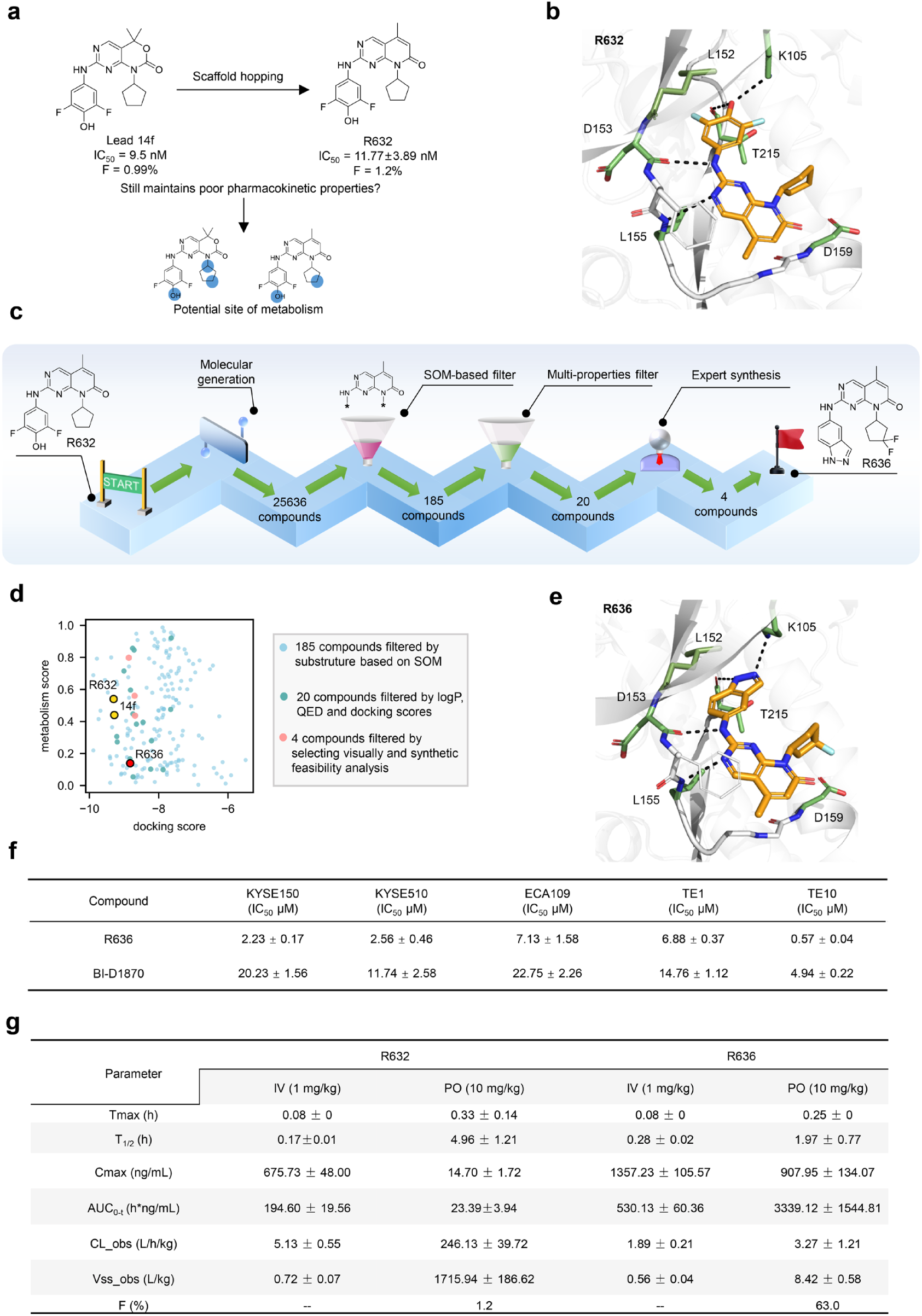
ConMeter is a key driver in the discovery of novel, potent and orally bioavailable RSK4 inhibitor R636. **a**, Identification of critical SOMs (indicated by blue circles) in lead compound 14f and R632. **b**, The predicted binding mode of R632 in the ATP-binding site of RSK4 NTKD. Crucial interacting residues are displayed as green sticks. Hydrogen bonds are depicted as black dashed lines. **c**, Rational optimization of R632 via molecular generation, SOM-driven substituent screening, multi-parameter prioritization yields high-priority candidates. d, The scatter diagram of metabolism scores and docking scores based on the filtered compounds, 14f, R632, R636. Metabolism score is defined as the top 1 prediction probability of every reactant to assess the possibility of metabolism. **e**, The predicted binding mode of R636 in the ATP-binding site of RSK4 NTKD. Crucial interacting residues are displayed as green sticks. Hydrogen bonds are depicted as black dashed lines. **f**, Cellular anti-proliferation assay for R636 and the control BI-D1870. **g**, Pharmacokinetic parameters of R632 and R636. Data are presented as mean±SD.

To overcome the limitations of traditional medicinal chemistry approaches and gain a more comprehensive and predictive understanding of the metabolic landscape of 14f and R632, we strategically deployed our newly developed metabolite prediction model, ConMeter. ConMeter was employed to identify and prioritize susceptible SOMs for both 14f and R632. The results showed that the difluorophenol and cyclopentane substituents consistently emerged as high-probability SOMs, ranking within the TOP 5 predictions for both 14f and R632 (**Fig. 4a and Supplementary Table 1**). This consistent finding strongly suggested that these two structural moieties were primary SOMs vulnerability and therefore warranted focused attention for subsequent structural modification efforts.

To rationally identify suitable substituents for effectively blocking the predicted SOMs, we employed a synergistic strategy that seamlessly integrated a well-established molecular generation approach (**Extended Data Fig. 5a**) with our ConMeter-guided insights. We initially pre-trained a robust molecular generation model using a Long Short-Term Memory (LSTM) algorithm, meticulously trained on an expansive dataset of 2,699 chemical entries^40^. This enabled the *de novo* generation of a vast and structurally diverse chemical library, comprising 30,000 SMILES strings, of which 25,636 were chemically valid. Given the structural compatibility of the pyridone pyrimidine fragment of R632 with the RSK4 binding pocket, including critical hydrogen bonding interactions with Leu155 and ASP153, and essential hydrophobic interaction with Gatekeeper Leu152 (**Fig. 4b**), we strategically prioritized the preservation of this key pyridone pyrimidine scaffold throughout the molecular generation process. Navigating the selection of truly promising candidates from a vast pool of 25,636 molecules presents a formidable challenge in traditional drug discovery. However, our innovative SOM-based screening approach dramatically streamlined this selection process, providing precise and actionable guidance for targeted compound modification. By strategically focusing on maintaining the pyridone pyrimidine scaffold while selectively modifying the predicted metabolic hotspots in other structural regions, we efficiently identified a focused subset of 185 compounds containing the desired target substructures (**Fig. 4c, 4d**). To further refine this candidate pool, we implemented a rigorous multi-parameter filtering cascade, incorporating metabolite substructure searching and stringent criteria based on calculated logP, drug-likeness (QED), and Glide docking scores ^41^. This multi-faceted filtering process effectively narrowed down the candidates to a highly prioritized set of 20 molecules for subsequent in-depth evaluation (**Supplementary Data 2**).

To visualize the chemical space distribution of the generated compounds selected via SOM-based filtering, we applied UMAP dimensionality reduction based on their Morgan fingerprint structural representations (**Extended Data Fig. 5b**). The resulting UMAP projection revealed that, while exhibiting broad distribution across the RSK4-active chemical space, the generated compounds remarkably maintained spatial proximity to known RSK4-active small molecules. This spatial distribution pattern elegantly suggests that our approach successfully generated compounds possessing significant structural diversity while concurrently preserving and potentially enhancing RSK4 activity. These results demonstrate the power and effectiveness of integrating SOM-based filtering with advanced molecular generation approaches for the targeted and efficient exploration of relevant chemical space.

To further validate our computational design strategy and identify lead optimization candidates, we prioritized compounds with favorable synthetic accessibility (**Fig. 4c)**. Following a comprehensive synthetic feasibility analysis, four representative compounds (R633, R634, R635 and R636,) were ultimately selected for synthesis (**Extended Data Fig. 6a**). Subsequently, their bioactivities were tested through *in vitro* enzyme inhibition assays. The results showed that R633, R634, R635 and R636 all exhibited potent RSK4 inhibitory activity, with IC50 values of 56.67±4.94 nM, 86.29±40.70 nM, 45.88±3.11 nM and 13.75±0.65 nM, respectively (**Extended Data Fig. 6b**). Notably, R636 emerged as the most potent inhibitor within this series, showcasing a significantly enhanced inhibitory potency compared to the other synthesized analogs.

Specifically, R636 replaced the difluorophenol group present in earlier analogs with an indazole moiety and substituted the cyclopentane group with two fluorine atoms. Intriguingly, ConMeter predictions indicated a significant enhancement in metabolic stability for R636 resulting from these targeted structural alterations (**Supplementary Table 1**). Complementary molecular docking studies further elucidated the favorable binding mode of R636, revealing its capacity to effectively penetrate the critical inner subpocket of RSK4, engaging key residues Leu152, Lys105, and Thr215, and forming hydrogen bonds with Lys105 (**Fig. 4e**). Furthermore, based on the docking results, advanced molecular dynamics (MD) simulations and binding affinity calculations provided deeper mechanistic insights (**Extended Data Fig. 7a**), demonstrating that R636 had higher solvation energy for the crucial residue ASP159 compared to R633. This finding suggested that the difluoro substitution improved the binding profile of R636 (**Supplementary Table 2 and Extended Data Fig. 7b**), contributing to its enhanced potency and potentially improved pharmacokinetic characteristics.

Consistent with its optimized design and predicted properties, *in vitro* cellular anti-proliferation assays unequivocally demonstrated that R636 displays remarkable and broad-spectrum bioactivity against a panel of ESCC cell lines (**Fig. 4f**). Notably, TE-10 cells exhibited exceptional sensitivity to R636, with an impressively low IC50 value of 0.57 μM. Furthermore, R636 effectively inhibited the invasion of TE10 cells in a dose-dependent manner (**Extended Data Fig. 8**). The IC50 value of R636 for suppressing TE10 cell invasion was 0.5812 μM (**Extended Data Fig. 8d**). Meanwhile, R636 dose-dependently inhibited colony formation (**Extended Data Fig. 9a** and **9c**) and promoted apoptosis in TE10 cells (**Extended Data Fig. 9b** and **9d**).

To comprehensively evaluate the *in vivo* potential of R636, we proceeded to assess its pharmacokinetic properties in SD rats. Following a single intravenous administration of R636, the compound showed a moderate clearance rate (1.89 L/h/kg) and a pharmacologically relevant half-life (0.28 h). Crucially, upon oral administration, R636 demonstrated significantly improved systemic exposure, with a substantially enhanced AUC0-t value of 3339.12 ng/mL*h. This translated into a remarkable oral bioavailability of 63% (**Fig. 4g**), a dramatic improvement compared to the negligible bioavailability of the lead compound 14f. Collectively, these compelling pharmacokinetic indicators strongly suggest that the metabolic properties of R636 have been successfully and substantially improved compared to 14f, a conclusion that is remarkably consistent with the metabolic probability predictions generated by ConMeter (**Supplementary Table 1**).

### R636 specifically targets the RSK4 signaling pathway and displays potent antitumor efficacy in ESCC PDX models

Given the excellent *in vitro* anticancer activity of R636 observed in the TE10 cell line, this cell line was selected for in-depth mechanistic investigations. TE10 cells were treated with different concentrations of R636, and the phosphorylation levels of RPS6 and GSK3*β*, which are typical downstream substrates of RSK4 signaling, were detected by Western blotting. The results showed that R636 potently inhibited the phosphorylation of RPS6 and GSK3*β* in a clear dose-dependent manner. Notably, at a concentration of 10 μM, R636 achieved near-complete suppression of RPS6 and GSk3*β* phosphorylation, which was comparable to the level of inhibition observed with the same concentration of BI-D1870 (**Fig. 5a**). To further explore the target specificity of R636 and rule out off-target effects on other RSK family members (RSK1-3), we detected the phosphorylation status of established downstream substrates of RSK1-3, including EIF4B and P27KIP1 of RSK1, creb and HSP27 of RSK2, and c-fos and HistoneH3 of RSK3 (**Fig. 5b-5d**). Remarkably, we observed that the addition of R636 did not affect the phosphorylation levels of RSK1-3 substrates, regardless of whether RSK4 was knocked down or not. Based on these findings, we concluded that R636 specifically targets the RSK4 signaling pathway.

**Fig. 5.**
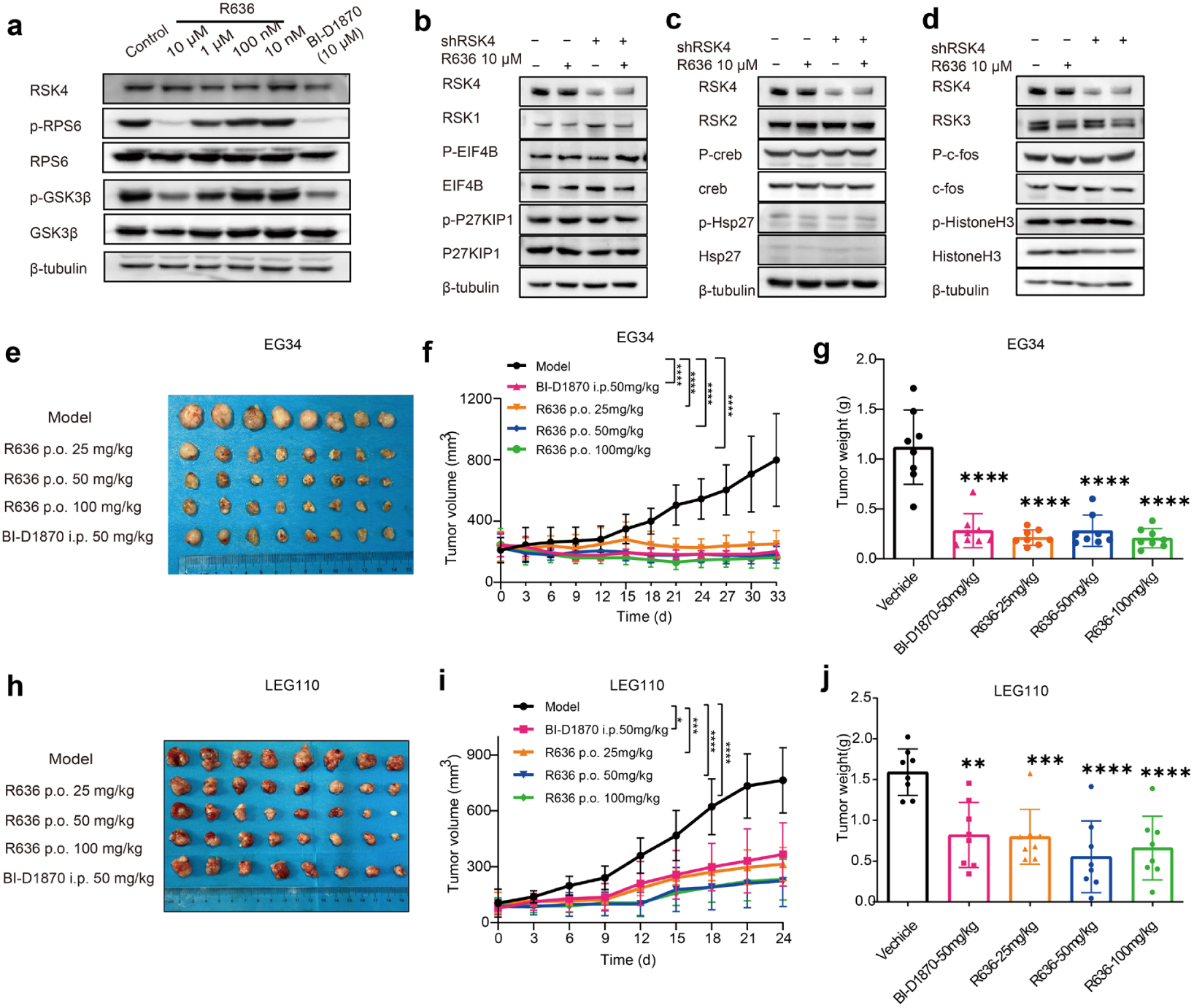
R636 specifically targets the RSK4 signaling pathway and demonstrates excellent anti-ESCC efficacy in two distinct PDX mouse models. **a**, Western analysis of the phosphorylation of RSK4 substrates with different doses of R636 for 24 h. BI-D1870 (10 μM) and DMSO were used as the positive and blank control. **b-d**, Western analysis of RSK1, RSK2, and RSK3 and the phosphorylation level of their corresponding substrates before and after the treatment of R636 (10 μM) for 24 h in the presence or absence of RSK4 knockdown. **e-g**, Tumor tissue picture of mice, tumor volumes change and tumor weight in EG34 PDX model. **h-j**, Tumor tissue picture of mice, tumor volumes change and tumor weight in LEG110 PDX model. Data are presented as mean±SD. One-way ANOVA test. (*p < 0.05, **p < 0.01, ***p < 0.001, ****p < 0.0001)

Next, compound R636 was subjected to acute and subacute toxicity evaluations in SD rats. In the acute toxicity test, rats were orally administered single doses of 200, 500 and 1000 mg/kg (**Extended Data Fig. 10a, 10b**). For the subacute toxicity experiment, animals received daily oral doses of R636 at 250, 500 and 1000 mg/kg continuously for 28 days (**Extended Data Fig. 10c, 10d**). Reassuringly, no mortality was observed in any treatment group throughout both acute and subacute toxicity evaluations following oral R636 administration. Furthermore, meticulous monitoring of the animals revealed no significant behavioral abnormalities or overt signs of toxicity during the experimental period, establishing an LD50 exceeding 1000 mg/kg. Histopathological examination via hematoxylin-eosin (HE) staining of major organs revealed no evidence of significant parenchymal injury or inflammatory cell infiltration (**Extended Data Fig. 10e**). Based on these comprehensive subacute toxicity data, the preliminary No Observed Adverse Effect Level (NOAEL) was conservatively estimated to be 250 mg/kg, with the Maximum Tolerated Dose (MTD) being greater than 1000 mg/kg, as no clinical adverse effects were observed at these dose levels. Collectively, these robust preclinical safety findings underscore that R636 exhibits a highly favorable safety profile *in vivo*, supporting its further development.

To investigate the *in vivo* antitumor efficacy of R636, we established two distinct patient-derived xenograft (PDX) mouse models, representing the heterogeneous nature of ESCC (**Fig. 5e-5g, 5h-5j**). R636 (25, 50, and 100 mg/kg) was orally administered into the PDX mice and BI-D1870 (50 mg/kg) was intraperitoneally injected into the mice as a positive control. Importantly, as depicted in **Extended Data Fig. 11a, 11b**, R636 treatment at all tested doses did not induce any discernible body weight loss in the treated animals, indicating good tolerability. Strikingly, in both PDX models, oral administration of R636 resulted in a profound and statistically significant inhibition of tumor growth (**Fig. 5e, 5h**), as evidenced by marked reductions in both tumor volume (**Fig. 5f, 5i**) and tumor weight (**Fig. 5g, 5j**). In the EG34 PDX model, oral administration of R636 at 100 mg/kg exhibited superior antitumor efficacy, achieving an impressive tumor growth inhibition (TGI) rate of 82.6%, surpassing the efficacy of intravenously administered BI-D1870 (TGI = 78.4%). In the LEG110 PDX model, oral administration of R636 at a dose of 50 mg/kg achieved a TGI rate of 70.8%, significantly outperforming intravenously administered BI-D1870 (TGI = 52.1%). These compelling *in vivo* efficacy data unequivocally demonstrate that orally administered R636 exerts a potent and clinically relevant inhibitory effect on ESCC tumor growth, exceeding the efficacy of a benchmark RSK4 inhibitor.

## Discussion

Upon entering the body, therapeutic agents encounter substantial stability challenges, which impose significant constraints on their chemical structures. Numerous compounds with potential pharmacological activity are often abandoned due to metabolic instability^42^. Consequently, the early and accurate identification of metabolically labile chemical groups is crucial to guide rational structural optimization during drug discovery, thereby accelerating development timelines and improving overall success rates^43^. However, the inherent complexities and high-throughput demands of *in vivo* drug metabolism assessments present substantial temporal and economic burdens for pharmaceutical research endeavors^1–3^. *In silico* drug metabolite prediction models offer a transformative solution, providing critical support for streamlining drug metabolism optimization by significantly reducing timelines and costs, enhancing prediction accuracy, and enabling efficient large-scale screening, thus establishing themselves as indispensable tools in contemporary drug development paradigms^42^. Nevertheless, limitations persist in existing metabolite prediction approaches. Previous research databases have failed to comprehensively cover the metabolic reactions of both endogenous and exogenous compounds. Additionally, there is a lack of standardized classification for metabolic reaction types, and existing algorithms have not fully exploited the intricate relationships between reactants and products, nor effectively leveraged the discriminatory power of both positive and negative training samples. As a result, AI-driven metabolite prediction models necessitate advancements in both reaction coverage and algorithmic optimization to achieve the high-precision predictions of metabolic properties demanded in modern drug discovery.

To tackle these challenges, we have meticulously constructed the most expansive and comprehensive human-specific drug metabolism database to date, characterized by its unprecedented breadth of reaction rule coverage. This database encompasses 11,665 experimentally validated human metabolic reactions, a scale unmatched by previously published resources. To overcome the inherent limitations of rule-based methods in scalability and generalizability, we employed a forward enumeration algorithm in conjunction with RDKit’s capabilities, resulting in 2,488 high-quality, non-redundant, and expertly validated metabolic reaction rules. This strategic combination of data scale and algorithmic refinement significantly expands the coverage of metabolic transformations addressable by our prediction framework, ensuring a robust and exceptionally comprehensive predictive capability.

Building on the aforementioned database, we developed ConMeter, a deep learning framework that synergistically integrates chemical feature interaction with contrastive learning algorithms. This innovative architecture enables ConMeter to effectively capture the intricate chemical nuances governing metabolic transformations. The framework was rigorously trained and validated using a five-fold cross-validation protocol, and its performance was optimized through a joint objective function incorporating both cross-entropy and contrastive loss components. During the prediction phase, the trained ConMeter outputs a finely-grained probability score for each reactant-metabolite pair, providing a quantitative measure of reaction likelihood and, critically, enabling the identification of SOMs for query molecules. Notably, ConMeter outperforms existing state-of-the-art models in predictive accuracy, achieving a substantial average improvement of approximately 20% in TOP5 accuracy, marking a significant advancement in the field.

To validate the application value of ConMeter within the drug discovery pipeline, we selected the therapeutically relevant target RSK4 for comprehensive experimental verification. To address the metabolic instability and low oral bioavailability of 14f and R632, ConMeter accurately identified their primary SOMs in difluorophenol and cyclopentane moieties, thereby providing valuable guidance for subsequent medicinal chemistry modifications. Furthermore, we employed a molecular generation algorithm to *de novo* design 25,636 valid compound structures and applied substructure filtering based on the identified SOMs, efficiently narrowing the selection to a focused set of 433 highly promising candidates. This intelligent combination of metabolic site prediction and substructure-based filtering dramatically reduced the candidate pool, saving both time and resources during the compound selection process. Leveraging a comprehensive multi-parameter ranking approach, incorporating drug-likeness properties (such as docking scores, QED, and logP), we meticulously analyzed the top 20 prioritized compounds and found that compounds like R636 exhibited both low predicted metabolic probability and favorable drug-like characteristics. As a result, we successfully executed targeted structural modifications, replacing the labile difluorophenol group with a metabolically more robust indazole moiety (R633: RSK4 IC50 = 56.67±4.94 nM) and substituting difluorocyclopentane for cyclopentane (R636: RSK4 IC50 = 13.75±0.65 nM). These rationally designed modifications were validated through *in vitro* kinase inhibition assays and pharmacokinetic studies. Remarkably, R636 exhibited a substantial improvement in oral bioavailability, reaching an impressive 63% in rats, representing a significant leap forward from the initial lead compound 14f. Furthermore, R636 demonstrated compelling anti-ESCC efficacy both *in vitro* and *in vivo*, particularly in two distinct PDX mouse models, where it effectively inhibited tumor growth through oral administration. The expedited discovery of R636 not only powerfully exemplifies the practical utility of ConMeter in accurately predicting and mitigating drug metabolic liabilities but also yields a promising orally bioavailable small molecule candidate for the treatment of esophageal squamous cell carcinoma.

The inherent metabolic instability of phenolic hydroxyl moieties stems from their predisposition to undergo hepatic enzyme-catalyzed phase II biotransformation processes, including sulfonation, glucuronidation, and methylation, which dramatically enhance urinary/hepatic clearance pathways and compromise therapeutic exposure, representing a fundamental pharmacokinetic constraint in medicinal chemistry optimization strategies targeting this functional group^35, 44–46^. Our study provides a viable strategy, exemplified by the ConMeter-guided design of R636, for overcoming the challenges associated with metabolically labile phenolic hydroxyl moieties in drug design.

Given its foundation on the most comprehensive human-specific drug metabolism database assembled to date, and coupled with its demonstrably high predictive accuracy, the ConMeter model possesses broad applicability for predicting the metabolic properties of small-molecule compounds targeting diverse therapeutic classes, including kinases, GPCRs, ion channels, and beyond. This wide-ranging utility has the potential to transform drug metabolism optimization, significantly enhancing the efficiency of lead optimization campaigns while simultaneously reducing the substantial economic and temporal costs associated with candidate drug discovery. In conclusion, by strategically leveraging ConMeter, we successfully guided the discovery of the orally bioavailable RSK4 inhibitor R636, underscoring its significant potential as a therapeutic candidate for ESCC and powerfully illustrating the transformative impact of artificial intelligence in both rational lead optimization and experimental validation within the pharmaceutical sciences.

## Methods

### Details of the metabolism database and template database

We compiled a comprehensive database of compound-metabolite pairs from several sources, including the MDL^47^ (version 2010) database, DrugBank^48^ (updated as of November 2, 2020), the biotransformation database (MetXBioDB)^49^ from BioTransformer, Human Metabolome Database (HMDB)^50^ (version 4.0), HumanCyc from MetaCyc^51^ (version 23.0), Recon3D^52^ (version 3.01), and the database from GLORYx. Through this process, we identified a total of 11665 reactions corresponding to 6932 unique reactants following the standardization of SMILES and protonation at pH7.5, while filtering out duplicate reactions. Subsequently, we categorized the reactions according to their reaction types.

Then, we defined the template extracting algorithm, which outlines a workflow for generating reaction rules. This algorithm begins by identifying the transformation region, characterized as the reaction center based on chemical properties such as hydrogens, charges, bonds, and rings. It then employs a generic atom as a wildcard to connect the transformation region. This algorithm constructs SMARTS-based templates for each metabolic reaction in the database, thereby facilitating the generation of a substantial collection of positive and negative samples using the RunReactant function of RDKit.

### Construction of the drug metabolite prediction model ConMeter

We have developed a machine learning framework to distinguish between true and false metabolites, leveraging chemical feature interaction and contrast learning algorithms for training. The process is structured as follows:

Each reactant and product is represented by a molecular fingerprint, which is constructed by concatenating three types of fingerprints: Extended Connectivity Fingerprint (ECFP)^53^, PubChem Fingerprint^54^, and ErG Fingerprint^55^. The resulting fingerprint vectors for reactants and products are each with 2346 dimensions.

Initially, we define the base model architecture as follows: we undergo dimensionality reduction through a shared preprocessing module consisting of a linear transformation layer (2346 → 1024 dimensions) and a GELU activation function. Then, the dimensionally reduced fingerprints are denoted as 𝑥_𝑟_ and 𝑥_𝑐_, respectively.

To further enhance the representation of reactants and products, we utilize two multi-layer perceptron (MLPs) to generate high-level feature embedding ℎ_𝑟_ = (W^(1)^ 𝑥_𝑟_)), and ℎ_𝑐_ = 𝜎(W^(2)^ ℎ_𝑐_)), where σ is the ReLU activation function. In the model, the vectors ℎ_𝑟_ and ℎ_𝑐_ are concatenated in two distinct sequences to form 1024-dimensional input vectors. Initially, ℎ_𝑟_ is concatenated with ℎ_𝑐_ and inputted into a neural network to yield the vector 𝑦_1_. Subsequently, the order is reversed, with ℎ_𝑐_ concatenated with ℎ_𝑟_ and this configuration is again processed through another network to produce the vector 𝑦_2_. The final predicted output, denoted as ŷ, is derived through a learnable weighted fusion mechanism. This mechanism integrates the outputs 𝑦_1_, 𝑦_2_, using dynamically adjustable coefficients 𝑎_1_, 𝑎_2_. These coefficients are initialized to ensure an unbiased contribution from each predictor and are implemented as trainable parameters within the neural network architecture. Throughout the training process, these coefficients are optimized via backpropagation, enabling the model to autonomously determine the optimal contribution weights for 𝑦_1_, 𝑦_2_ based on their respective significance to the prediction task.

This architecture provides a streamlined approach that focuses on fundamental feature extraction and adaptive fusion, serving as a baseline for comparing more complex models that incorporate the Chemical Feature Interaction Module and the Constractive Learning Module.

Then, we trained the base model with the Chemical Feature Interaction Module, the interaction between 𝑥_𝑟_ and 𝑥_𝑐_ is captured by the outer product to form a sparse matrix X_1_ = 𝑥_𝑟_×𝑥*_c_*. This matrix is then transformed by multiplying two variable matrices into a smaller matrix X_𝟐_, which is subsequently reshaped into a 1024-dimensional vector 𝑥_𝟑𝟑_. The vector 𝑥_𝟑𝟑_ is fed to a four-layer fully connected neural network to produce an output referred to as an one-dimension vector 𝑦_3_. The same adaptive fusion mechanism is applied to get ŷ through the dynamic weighted aggregation among 𝑦_1_, 𝑦_2_ and 𝑦_3_. Furthermore, the base model with the Contrastive Learning Module is constructed. ℎ_𝑟_and ℎ_𝑐_ in base model are further transformed into latent vectors respectively, 𝑧_𝑟_ = 𝜎(W^(3)^ ℎ_𝑟_)**, and** 𝑧_𝑐_ = 𝜎(W^(4)^ ℎ_𝑐_).

To distinguish positive reactant-product pairs from negative pairs, a contrastive loss function is defined using 𝑧_𝑟_ and 𝑧_𝑐_, ensuring that positive pairs are brought closer in the embedding space, while negative pairs are pushed apart.

Let sim(𝑢,𝑣) = 𝑢^T^𝑣⁄‖𝑢‖‖𝑣‖ denote the cosine similarity between l2 normalized vectors ***u*** and ***v***. The loss function of positive and negative pairs from a single reactant is defined as:

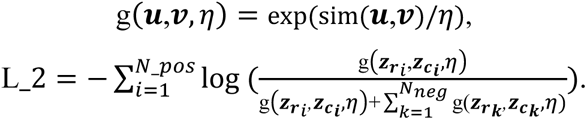

Here, *N_pos* indicates the number of positive pairs while N*_neg* indicates the number of negative pairs.

During training process, the optimization is performed using a combined loss function comprising cross-entropy loss, denoted as L_1 and contrastive loss L_2.

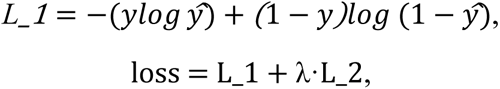

where λ is a constant defined as λ = 0.1.

Finally, to enhance model performance and robustness, we implemented an ensemble approach through the same adaptive fusion mechanism. This architecture integrates four pretrained models: The model with Chemical Feature Interaction Module and the model with Contrastive Learning Module, pretrained with ROC-AUC optimization and PRC-AUC optimization, respectively.

### Details of the evaluation on the test datasets compared with existing tools

To assess the predictive capabilities of ConMeter, we conducted a comparative analysis of four existing drug metabolite prediction tools: GLORYx^20^, BioTransformer^19^, MetaTrans^21^, and SyGMa^18^, to predict metabolites through a single-step reaction assessment. GLORYx and SyGMa employ rule-based approaches focused on oxidative and transferase reactions, specifically targeting phase I and II xenobiotic metabolism. In contrast, MetaTrans and BioTransformer offer broader enzymatic coverage, with BioTransformer incorporating an “allHuman” module encompassing transformations mediated by both human enzymes and gut microbiota.

To compare the performance of MetaTrans, GLORYx, and SyGMa, we ranked the top 5, 10, 13, and 20 predicted metabolites against the entire set of metabolites. This ranking also aligns with the beam size specified by MetaTrans to control output size. Additionally, we chose the top 13 to facilitate a fair comparison with BioTransformer, whose specific test set yields an average output size of 13. We used the Inchi representation to evaluate all methods.

### Molecular generation model for potential RSK4 activities

A molecular generation model based on Long Short-Term Memory (LSTM) architecture was pre-trained using RPS6 kinase (UniProt: Q9UK32)^56^ active compounds from ChEMBL (ID: CHEMBL4924)^57^. Initial curation removed 394 inactive molecules and duplicates, resulting in 248 unique compounds via RDKit^37^. Data augmentation expanded the dataset to 2,699 entries. Then the dataset was partitioned into training (n=2,154) and testing (n=545) sets. The model underwent 20-epoch pre-training followed by 5-epoch fine-tuning using lead compound R632.

### Molecule docking

The compounds were prepared utilizing the LigPrep module employing the OPLS_3 force field. A total of 32 stereoisomers were generated for each compound. The RSK4 crystal structure (PDB code: 6G77) was prepared using Protein Preparation Wizard available in Maestro, with docking grids centered on the co-crystallized ligand centroid. Then, the Induced Fit Docking was performed using Schrödinger Maestro (v10.1). We checked the Trim side chains option, and selected the gatekeeper residue Leu152. Glide was employed for docking simulation in standard precision (SP) mode. Next, using the complex of 14f and RSK4 as the reference structure, the other derivatives were docked with RSK4 by the Ligand Docking module in Maestro.

The docking scores are in **Supplementary Data 2**. The complexes of compound with protein RSK4 are saved in the format of PDB files, also as the initial conformation for molecular dynamics simulations.

### Molecular dynamics simulations

AMBER14 with ff99SB force field was implemented for simulations of the complexes of R632, R633, and R636 with protein RSK4. Ligand parameters were generated using Antechamber, with systems solvated in TIP3P water boxes and neutralized by counterions. The systems were subjected to descending restraint forces for minimization. Subsequently, the temperature was gradually increased from 0 to 300 K under NVT ensemble conditions, followed by equilibration at 300 K under NPT ensemble conditions for a duration of 500 ps. Finally, each system underwent an unrestrained MD production run lasting for 100 ns. Additionally, we estimated the binding energy of the complexes using MM/GB(PB)SA methods implemented in the mmpbsa.py tool of AmberTools18 by analyzing snapshots from each system over a span of 50 instances.

### Chemical synthesis

#### Synthesis and characterization of R636, R632, R633, R634 and R635. 5-bromo-2-chloro-*N*-(3,3-difluorocyclopentyl)pyrimidin-4-amine (2)

To a solution of 5-bromo-2,4-dichloropyrimidine (0.20 g, 0.88 mmol) in 5 mL acetonitrile, potassium carbonate (0.22 g, 1.58 mmol) was added at 0 °C. Then, a solution of 3,3-difluorocyclopentan-1-amine (0.13 g, 0.85 mmol) in 1 mL of acetonitrile was added dropwise. The mixture was stirred at room temperature for 6 h. After completion, ice water was added, and the mixture was extracted with dichloromethane. The organic layer was washed with saturated aqueous sodium chloride, dried over anhydrous sodium sulfate, and concentrated in vacuo. Purification was carried out by silica gel chromatography (dichloromethane/methanol = 200:1, v/v) to give product 5-bromo-2-chloro-*N*-(3,3-difluorocyclopentyl)pyrimidin-4-amine as white solid. LC-MS: m/z: 312.0 (M+H)^+^.

#### 2-chloro-8-(3,3-difluorocyclopentyl)-5-methylpyrido[2,3-*d*]pyrimidin-7(8*H*)-one (3)

To a solution of 5-bromo-2-chloro-*N*-(3,3-difluorocyclopentyl)pyrimidin-4-amine (0.10 g, 0.32 mmol) in 25 mL three-necked flask, palladium acetate (7.20 mg, 0.30 mmol), crotonic acid (42 mg, 0.48 mmol) and triethylamine (130 mg, 1.29 mmol) were added. The flask was purged with nitrogen three times, and then 5 mL of *N*-methylpyrrolidone was added. The reaction mixture was heated at 75 °C, and thin-layer chromatography (TLC) analysis indicated that the reaction was complete. Then, acetic anhydride (0.2 mL) was added, and the mixture was further heated at 75 °C until full conversion to the desired product was achieved. After cooling, the reaction mixture was extracted with dichloromethane and water. The combined organic layers were dried over anhydrous sodium sulfate and concentrated under reduced pressure. The crude product was subjected to 300-400 mesh silica gel column chromatography (petroleum ether/ethyl acetate=10/1, v/v) to obtain 65 mg of yellow solid. ^1^H NMR (400 MHz, DMSO-*d6*) δ 8.29 (s, 1H), 7.72 (s, 1H), 4.63-4.49 (m, *J* = 7.1 Hz, 1H), 2.51 (s, 3H), 2.44 – 2.19 (m, 3H), 2.17-2.10 (m, 2H), 2.00 – 1.86 (m, 1H). LC-MS: m/z: 312.0 (M+H)^+^.

#### 2-((1*H*-indazol-5-yl)amino)-8-(3,3-difluorocyclopentyl)-5-methylpyrido[2,3- *d*]pyrimidin-7(8*H*)-one (R636)

To a solution of 2-chloro-8-(3,3-difluorocyclopentyl)-5-methylpyrido[2,3- *d*]pyrimidin-7(8*H*)-one (0.05 g, 0.17 mmol) and *p*-toluenesulfonic acid (0.03 g, 0.19 mmol) in 1mL of *n*-butanol, 1*H*-indazol-5-amine (0.03 g, 0.20 mmol) was added. The mixture was heated for 1h at 100 °C, and TLC analysis indicated that the reaction was complete. After cooling, the reaction mixture was extracted with dichloromethane and aqueous sodium bicarbonate solution. The combined organic layers were dried over anhydrous sodium sulfate and concentrated under reduced pressure. Purification was carried out by slica gel chromatography (dichloromethane/methanol = 20:1, v/v) to give **(R636)** as yellow solid ( 0.03 g, 44.78%).^1^H NMR (600 MHz, DMSO-*d*6) *δ* (ppm): 13.00 (s, 1H), 10.05 (s, 1H), 8.85 (s, 1H), 8.17-8.12 (m, 1H), 7.97 (s, 1H), 7.56 (dd, *J* = 8.9, 1.9 Hz, 1H), 7.50 (d, *J* = 8.9 Hz, 1H), 6.21 (d, *J* = 1.4 Hz, 1H), 6.11 (s, 1H), 3.03 (dq, *J* = 27.6, 11.9 Hz, 1H), 2.49-2.41 (m, 2H), 2.39 (d, *J* = 1.2 Hz, 3H), 2.28 (ddt, *J* = 19.0, 13.2, 6.5 Hz, 1H), 2.12-1.96 (m, 2H). ^13^C NMR (151 MHz, DMSO-*d*6) *δ* 163.11, 159.66, 157.58, 155.76, 146.57, 137.31, 133.61, 132.85, 130.12, 123.33, 122.35, 117.42, 111.07, 110.48, 49.79, 36.51, 34.22, 29.50, 27.02, 17.22. HRMS (ESI) (m/z): [M+H]^+^ calcd for C20H18F2N6O, 397.1589; found 397.1589.

#### 8-Cyclopentyl-2-((3,5-difluoro-4-hydroxyphenyl)amino)-5-methylpyrido[2,3- *d*]pyrimidin-7(8*H*)-one (R632)

m.p 244.3-245.1 °C。^1^H NMR (400 MHz, DMSO-*d*6) δ 10.01 (s, 1H), 9.77 (s, 1H), 8.83 (s, 1H), 7.44 (d, *J* = 10.0 Hz, 2H), 6.21 (s, 1H), 5.89-5.69 (m, 1H), 2.37 (s, 3H), 2.25-2.20 (m, 2H), 1.93-1.80 (m, 2H), 1.83-1.69 (m, 2H), 1.67-1.53 (m, 2H). ^13^C NMR (151 MHz, DMSO-*d*6) δ 162.88, 158.92, 157.23, 155.94, 152.55 (d, *J* = 238.7 Hz), 152.49 (d, *J* = 239.0 Hz), 145.77, 131.70, 129.02, 118.33, 107.66, 104.04, 103.99, 53.35, 27.93, 25.45, 17.13. HRMS (ESI) (m/z): [M+H]^+^ calcd for C19H18F2N4O2, 373.1476; found 373.1475.

#### 2-((1H-indazole-5-yl)amino)-8-cyclopentyl-5-methylpyridine[2,3-*d*]pyrimidine-7 (8*H*)-one (R633)

m.p 238.3-239.1 °C。^1^H NMR (400 MHz, DMSO-*d6*) δ 13.00 (s, 1H), 9.94 (s, 1H), 8.81 (s, 1H), 8.12 (s, 1H), 8.00 (s, 1H), 7.53 (q, *J* = 8.7 Hz, 2H), 6.18 (s, 1H), 5.84-5.80 (m, 1H), 2.37 (s, 3H), 2.25-2.20 (m, 2H), 1.88-1.69 (m, 4H), 1.56-1.50 (m, 2H). ^13^C NMR (151 MHz, DMSO-*d6*) δ 163.01, 159.72, 157.27, 156.04, 145.83, 137.32, 133.56, 132.98, 123.33, 122.52, 117.61, 111.20, 110.42, 107.20, 53.01, 27.95, 25.45, 17.15. HRMS (ESI) (m/z): [M+H]^+^ calcd for C20H20N6O, 361.1780; found 361.1779.

#### 8-(3,3-difluorocyclopentyl)-5-methyl-2-((2-oxoindol-5-yl)amino)pyrido[2,3- *d*]pyrimidin-7(8*H*)-one (R634)

m.p 259.4-259.9 °C。^1^H NMR (400 MHz, DMSO-*d6*) δ 13.03 (s, 1H), 10.04 (s, 1H), 8.82 (d, *J* = 22.3 Hz, 1H), 7.98 (s, 1H), 7.53 (d, *J* = 8.3 Hz, 2H), 6.19 (d, *J* = 22.3 Hz, 1H), 5.39-5.33 (m, 1H), 2.37 (s, 3H), 1.88-1.23(m, 10H). HRMS (ESI) (m/z): [M+H]^+^ calcd for C21H22N6O, 375.1933; found 375.1932.

#### 2-((1*H*-indol-5-yl)amino)-8-cyclohexyl-5-methylpyrido[2,3-*d*]pyrimidin-7(8*H*)- one (R635)

m.p 284.7-285.6 °C。^1^H NMR (400 MHz, DMSO-*d6*) δ 10.33 (s, 1H), 9.93 (s, 1H), 8.81 (s, 1H), 7.60 (s, 1H), 7.42 (d, *J* = 8.2 Hz, 1H), 6.77 (d, *J* = 8.0 Hz, 1H), 6.17 (d, *J* = 16.0 Hz, 1H), 6.15-6.00 (m, 1H), 3.46 (s, 2H), 3.01 (td, *J* = 24.0, 11.9 Hz, 1H), 2.49-2.40 (m, 2H), 2.37 (s, 3H), 2.29-2.19 (m, 1H), 2.19-2.00 (m, 2H). ^13^C NMR (151 MHz, DMSO-*d6*) δ 176.69, 163.09, 159.47, 157.52, 155.72, 146.53, 139.67, 133.88, 126.46, 120.40, 118.25, 117.35, 109.27, 60.83, 36.48, 35.16, 19.11, 17.19, 14.32. HRMS (ESI) (m/z): [M+H]^+^ calcd for C21H19F2N5O2, 412.1584; found 412.1584.

### Cell culture

The human ESCC cell lines TE10 and KYSE-510 were purchased from Nanjing Cobioer Biotechnology Co., Ltd. The human ESCC cell lines KYSE-150, TE1 and ECA-109 were purchased from the Type Culture Collection of the Chinese Academy of Sciences. All cell lines were cultured in RPMI 1640 medium supplemented 10% fetal bovine serum (Gibco, USA) and cultured at 37 ℃ in a humidified incubator with 5% CO2.

### In vitro enzymatic activity assay

All enzymatic reactions were performed at 30 °C for 40 minutes in 50 μL reaction systems containing: 40 mM Tris-HCl (pH 7.4), 10 mM MgCl₂, 0.1 mg/mL BSA, 1 mM DTT, 10 μM ATP, 0.2 μg/mL kinase, and 100 μM lipid substrate. Compounds were dissolved in 10% DMSO, and 5 μL of the solution was added to each 50 μL reaction mixture, resulting in a final DMSO concentration of 1% in all reactions. Kinase activity was measured using the Kinase-Glo Plus Luminescence Kinase Assay Kit, which quantifies the remaining ATP in the solution after the kinase reaction. IC₅₀ values were calculated using Prism GraphPad software.

### Cell viability assay

Cell viability was assessed using the CCK-8 assay kit (Beyotime). Cells were seeded in 96-well plates at 3000 cells/well in 100 μL RPMI 1640 medium. Subsequently, cells were treated with different concentrations of compounds and incubated for 72 hours. Then, 10 μL of the CCK-8 reagent was added to each well and plates were incubated at 37 °C for 2 h in the dark. Absorbance was measured at 450 nm. The IC50 values were calculated by using GraphPad Prism.

### Western blotting

Cells (6 × 10^6^ cells per well) were seeded in 6-well plates, incubated overnight, and treated with different concentrations of compounds for 24h. Harvested cells were washed twice using ice-cold PBS. Proteins were extracted using RIPA lysis buffer and separated via 10% SDS-PAGE, then transferred onto PVDF membranes. Membranes were blocked with 5% BSA for 2 hours at room temperature, the membranes were subsequently incubated with primary antibodies at 4 ℃ overnight. Immunoreactive bands were detected using species-matched horseradish peroxidase (HRP)-conjugated secondary antibodies at room temperature. The primary antibody include RSK4, p-RSK4(Ser232), GSK3β, p-GSK3β(Ser9), RPS6, p-RPS6(Ser235/236), β-tubulin, Caspase-3, cleaved-Caspase-3, PARP, cleaved-PARP and Bcl-2.

### Pharmacokinetic properties

The pharmacokinetic experiments were conducted by HangzhouLeading Pharmatech Co., Ltd. SD male rats were purchased from Shanghai SLAC Laboratory Animal Co., Ltd. The solvent used for drug administration consisted of 5% DMSO, 5% 100% Solutol, and 90% physiological saline. The doses were administered intravenously (IV) at 1 mg/kg and via oral gavage (IG) at 10 mg/kg. Liquid chromatography-tandem mass spectrometry (LC-MS/MS) analysis was conducted using an AB SCIEX Qtrap 5500 ion trap mass spectrometer, with data analysis performed using WinNonLin 8.0 software.

### Acute toxicity and subacute toxicity

The subacute toxicity experiments were conducted by HangzhouLeading Pharmatech Co., Ltd. SD rats were utilized to assess the safety of the test compound, with the animals randomly assigned to different dosage groups. Observations were conducted for 24 hours to 28 days post-administration, monitoring clinical signs, body weight changes, and mortality rates. After the study, the animals underwent necropsy and histopathological examination to evaluate potential chronic toxicity and organ damage.

### ESCC PDX models

This study was approved by the Research Ethics Committee of Zhengzhou University. All animal procedures in this study were approved by the Zhengzhou University Institutional Animal Care and Use Committee (ZZUIRB2022-72). All mice were fed with free access to food and water and kept in 12 h/12 h light/dark cycles. For ESCC PDX models, the tumor was taken after surgical operation and implanted into the NPG (Vital River Labs, Beijing, China) mice under a sterile environment. The measurement of tumor size was performed twice a week using vernier caliper and the calculation of tumor volume was done according to tumor volume (mm^3^)=length× width×height×0.52.

### Statistics analysis

Statistical analysis was conducted using GraphPad Prism software and the results were presented as mean ± SD. For multiple comparisons, an ordinary one-way analysis of variance (ANOVA) was used. A P-value of < 0.05 was considered a statistically significant difference.

### Code availability

The source code of ConMeter and the metabolism database can be downloaded from the GitHub repository at https://github.com/mzz1503/ConMeter

## Acknowledgement

This work was supported in part by the National Key Research and Development Program of China (2022YFC3400504, 2022YFC3400501); the National Natural Science Foundation of China (82425104 to H.L.; 82173690 to S.L.); the Shanghai Rising-Star Program (23QA1402800 to S.L.); the Key Research and Development Program of Shaanxi Province (2024SF-ZDCYL-02-05 to Z.W.); the Advanced Pathology Technology Research and Innovation Platform of the Shaanxi Provincial Health Commission Program (2023PT-04 to Z.W.); the State Key Laboratory of Holistic Integrative Management of Gastrointestinal Cancers: Self-initiated Research Projects (2025GTKP003 to Z.W.).

## Author contributions

H.H., M.Z., X.Y., and S.H. contributed equally to this work. S.L., K.L., K.Z., Z.W., and H.L. conceived and codirected the study. H.H. performed compound synthesis with advice from S.L., H.L., Z.Z., and X.Q.. M.Z. constructed the AI model and performed the computational simulation studies with advice from S.L., K.Z., and H.L.. C.L. N.C., T.Y., L.Z., X.Z., R.Z. and R.W. performed the toxicity experiment. F.H. performed molecular generation. X.S., X.Z., and Z.C. performed the evaluation of cell viability. S.H., Y.Y. and Z.W. performed the experiments on anti-invasion and cell signaling pathways. X.Y., K.L. conducted the in vivo PDX experiment. H.H., M.Z., prepared figures and wrote the manuscript. S.L., K.L., K.Z., Z.W. revised the manuscript. All authors have given approval to the final version of the manuscript.

## Competing Interests

The authors declare no competing interests.

## Extended Data Figures

**Extended Data Fig. 1.**
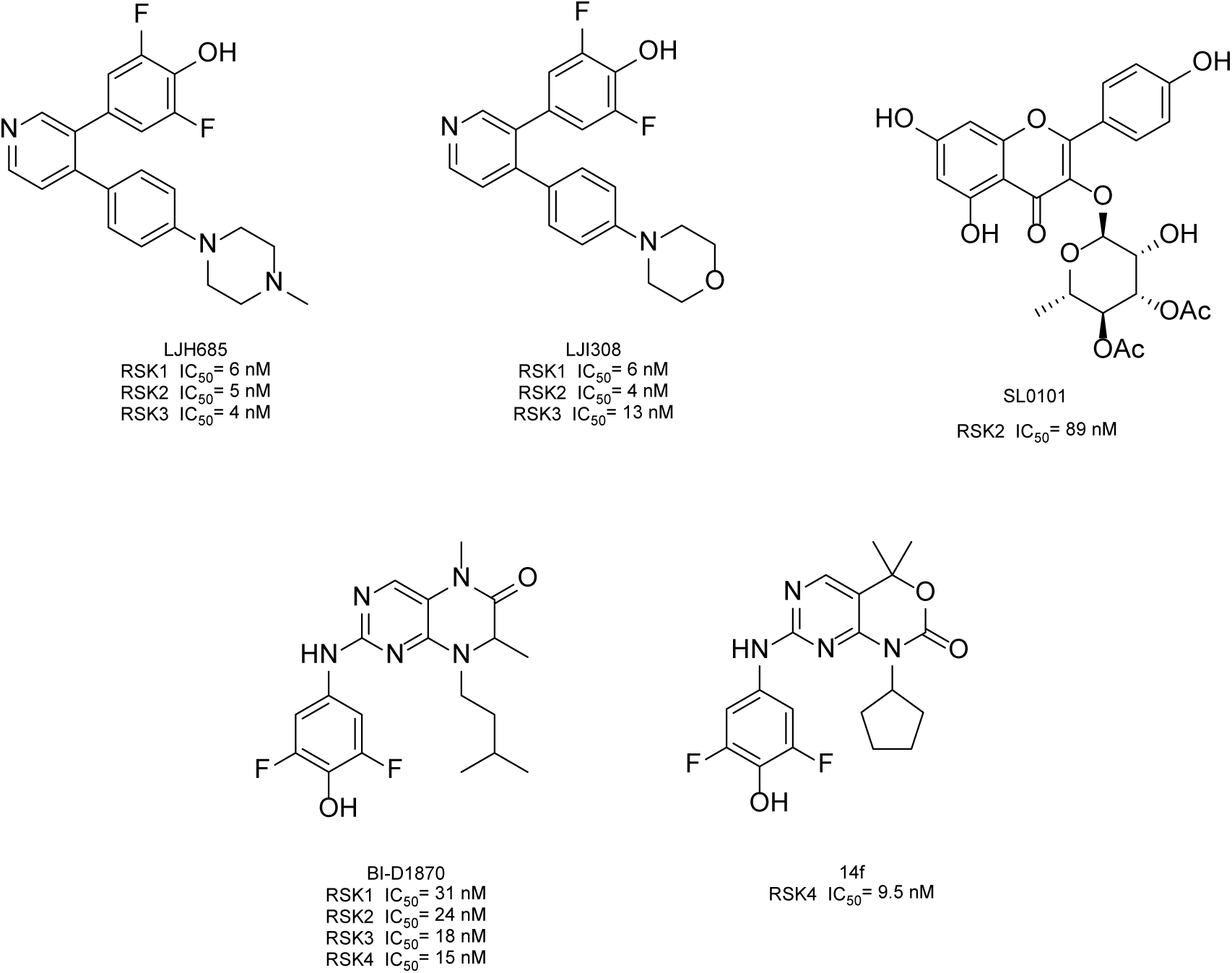
Chemical structures of representative RSKs inhibitors.

**Extended Data Fig. 2.**
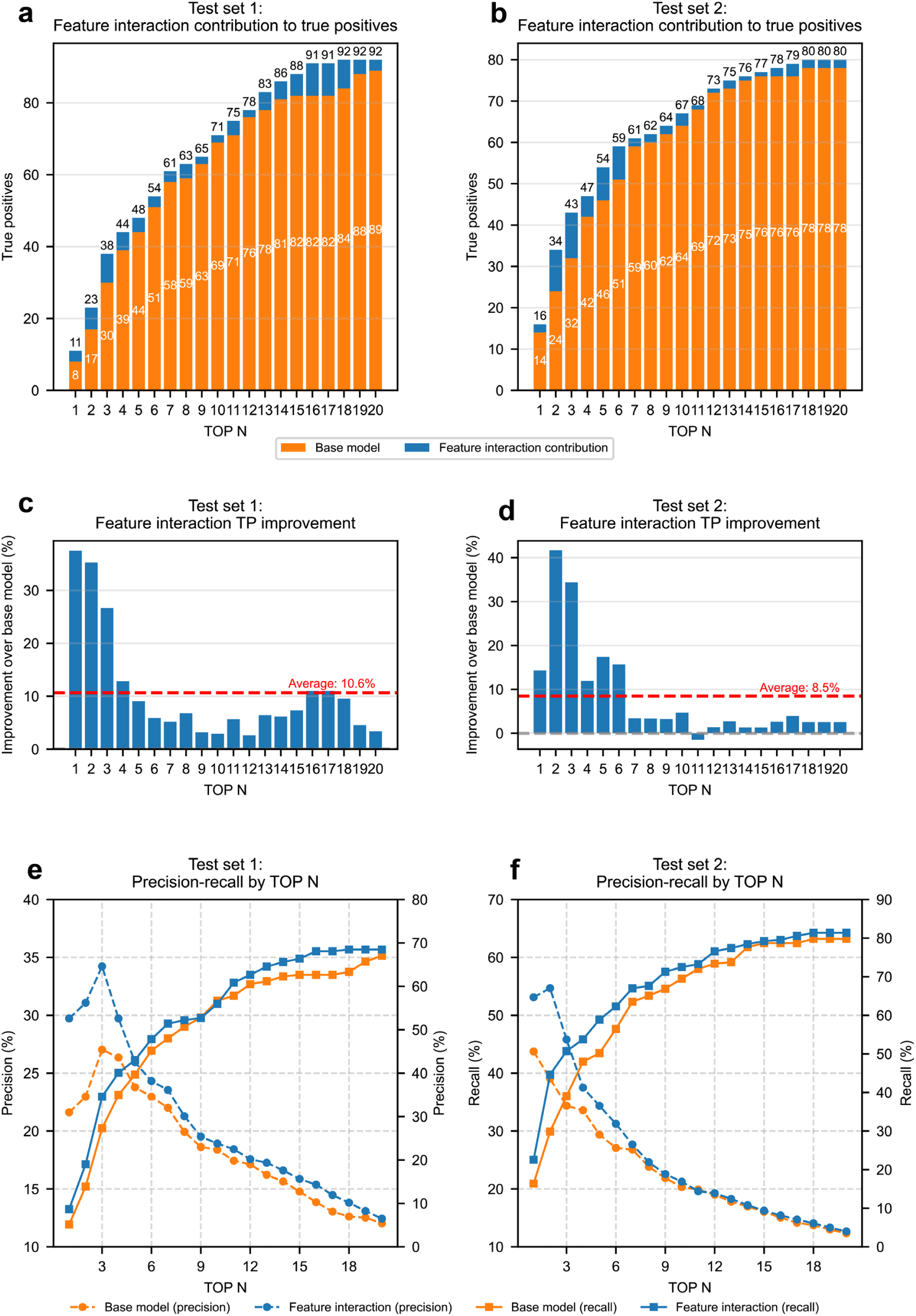
Performance evaluation of the Chemical Feature Interaction Module. **a-b**, Contribution to true positives compared to the base model across different TOP N values for Test Sets 1 and 2. **c-d**, Percentage improvement in true positive (TP) identification over the base model, with average improvements of 10.6% and 8.5% for Test Sets 1 and 2, respectively. **e-f**, Precision-recall curves demonstrating enhanced precision at lower TOP N values and consistent recall improvements across the full TOP N range.

**Extended Data Fig. 3.**
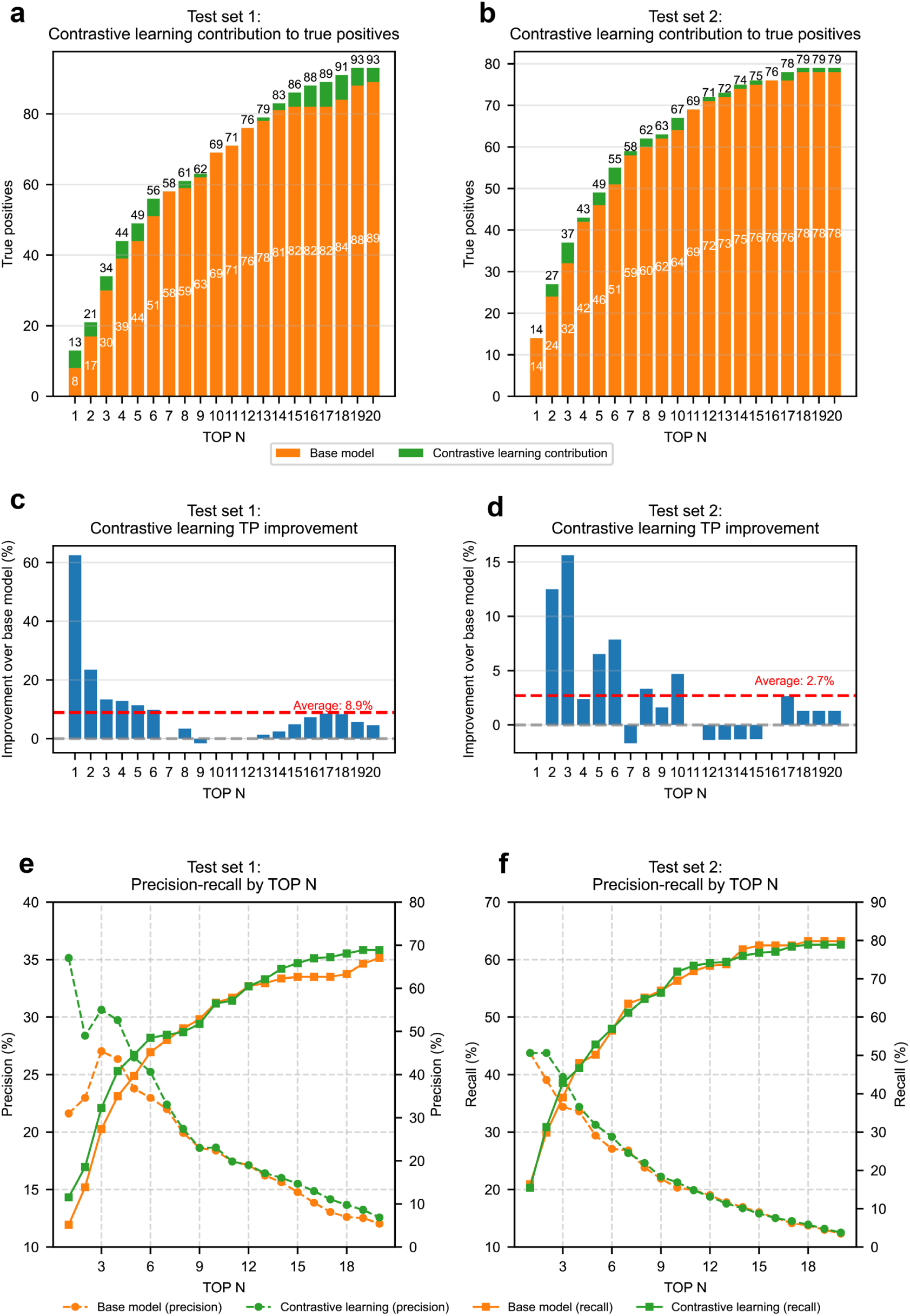
Performance evaluation of the Contrastive Learning Module. **a-b**, Contribution to true positives across different TOP N values for Test Sets 1 and 2, showing significant enhancements over the base model. **c-d**, Percentage improvement in true positive identification, with exceptional gains of up to 61% at TOP N = 1 in Test Set 1 and average improvements of 8.9% and 2.7% across both test sets. **e-f**, Precision-recall analysis illustrating the module’s effectiveness in maintaining higher precision at critical lower TOP N values while improving recall performance.

**Extended Data Fig. 4.**
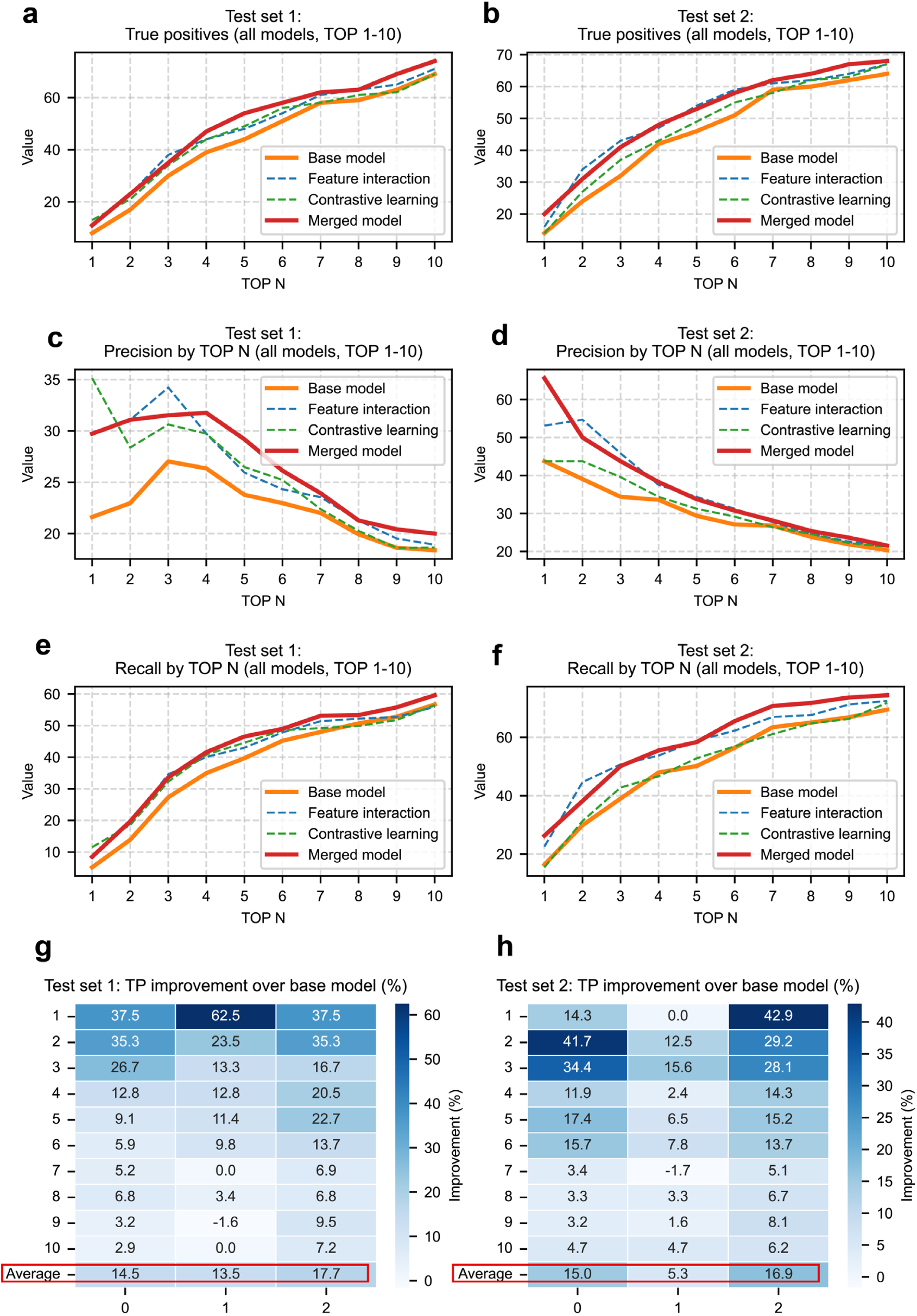
Comparative performance analysis of the merged model with Chemical Feature Interaction model, Contrastive Learning model and base model. **a-b**, True positive identification for TOP N values 1-10 across both test sets, demonstrating the merged model’s superior performance. **c-d**, Precision comparison showing the merged model’s balanced optimization of precision metrics. **e-f**, Recall analysis illustrating consistent improvements over the base model across all TOP N values. **g-h**, Heatmaps quantifying percentage improvements over the base model, with the merged model achieving the highest average improvements of 17.7% and 16.9% for Test Sets 1 and 2, significantly outperforming individual modules.

**Extended Data Fig. 5.**
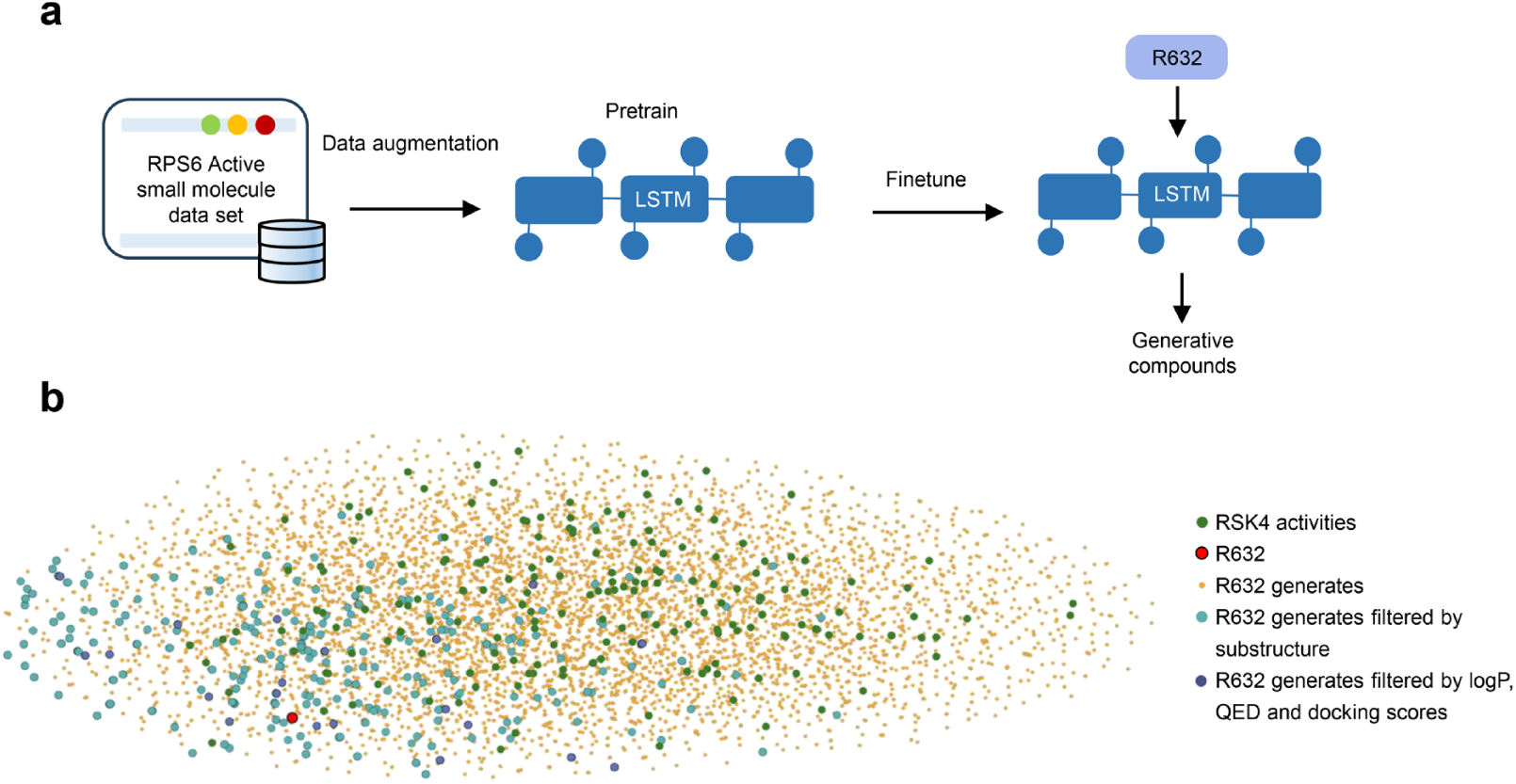
Details of the generation model. **a**, The training process of the molecule generation model. **b**, A U-MAP visualization of the result for molecular generation. The figure includes several categories of molecules: RSK4 activity molecules curated from the ChEMBL database (green), the R632 molecule (red), generated molecules based on R632 from the LSTM model (yellow), and molecules filtered by SOM-based substructure matching (blue). This visualization facilitates an intuitive understanding of the relationships between different molecules and their distribution within the feature space.

**Extended Data Fig. 6.**
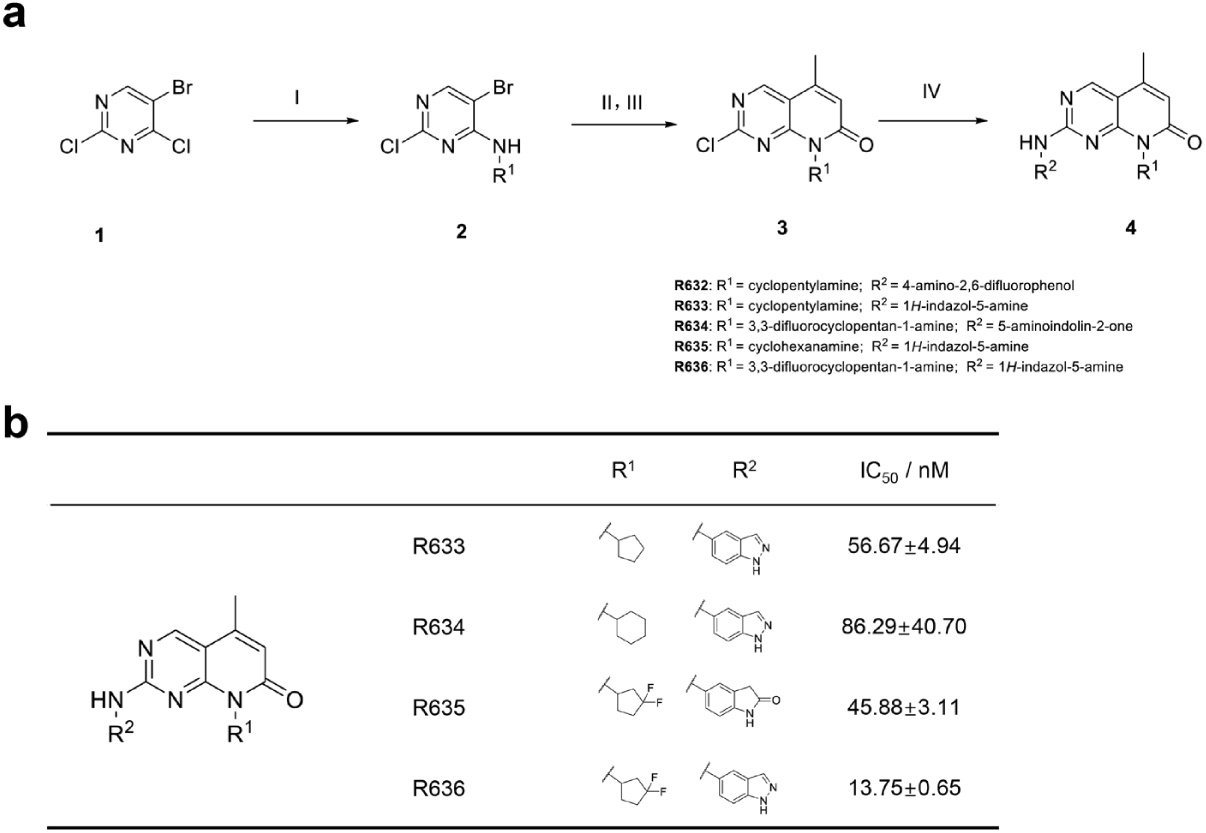
The synthesis and inhibitory activities of R632, R633, R634, R635 and R636. **a**, Synthesis of R632, R633, R634, R635 and R636. Reagents and conditions: (Ⅰ) R-NH2, K2CO3, MeCN, r.t.; (Ⅱ) Crotonic acid, (C6H5CN)2PdCl2, DIPEA, Tri(o-tolyl)phosphine, *n*-butanol, 95 °C; (Ⅲ) (CH3CO)2O, 85 °C; (Ⅳ) TFA, 2-butanol, ArNH2, 110 °C. **b**, The synthesized compounds R633, R634, R635 and R636 and their corresponding RSK4 enzyme activity data. Inhibitory activities are presented as the mean ± SD.

**Extended Data Fig. 7.**
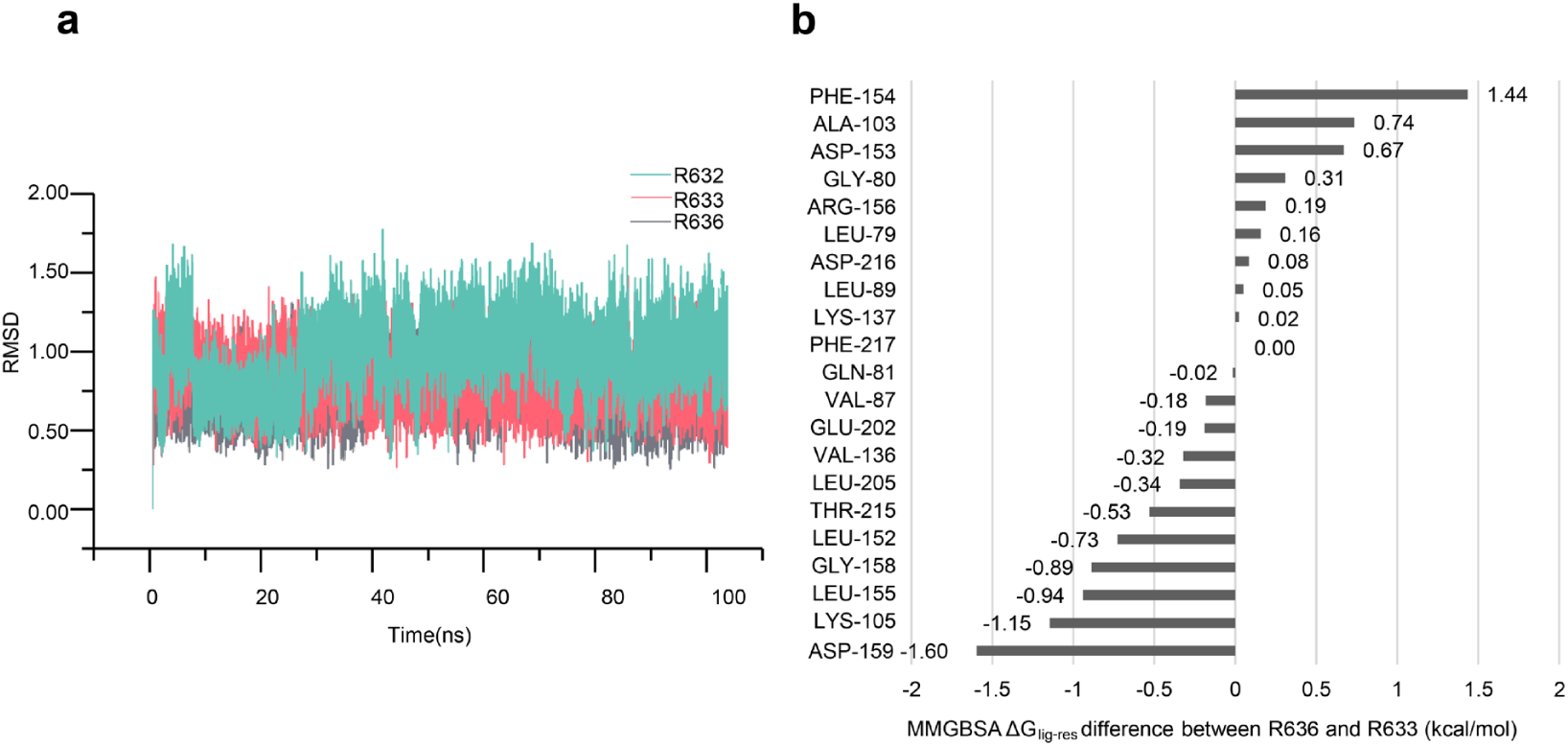
Results of molecular dynamics simulations. **a**, The RMSD plots of the ligands R632, R633, R636 for 100 ns MD simulations. **b**, MM-GBSA residue–ligand binding free energy decomposition analysis between R636 and R633.

**Extended Data Fig. 8.**
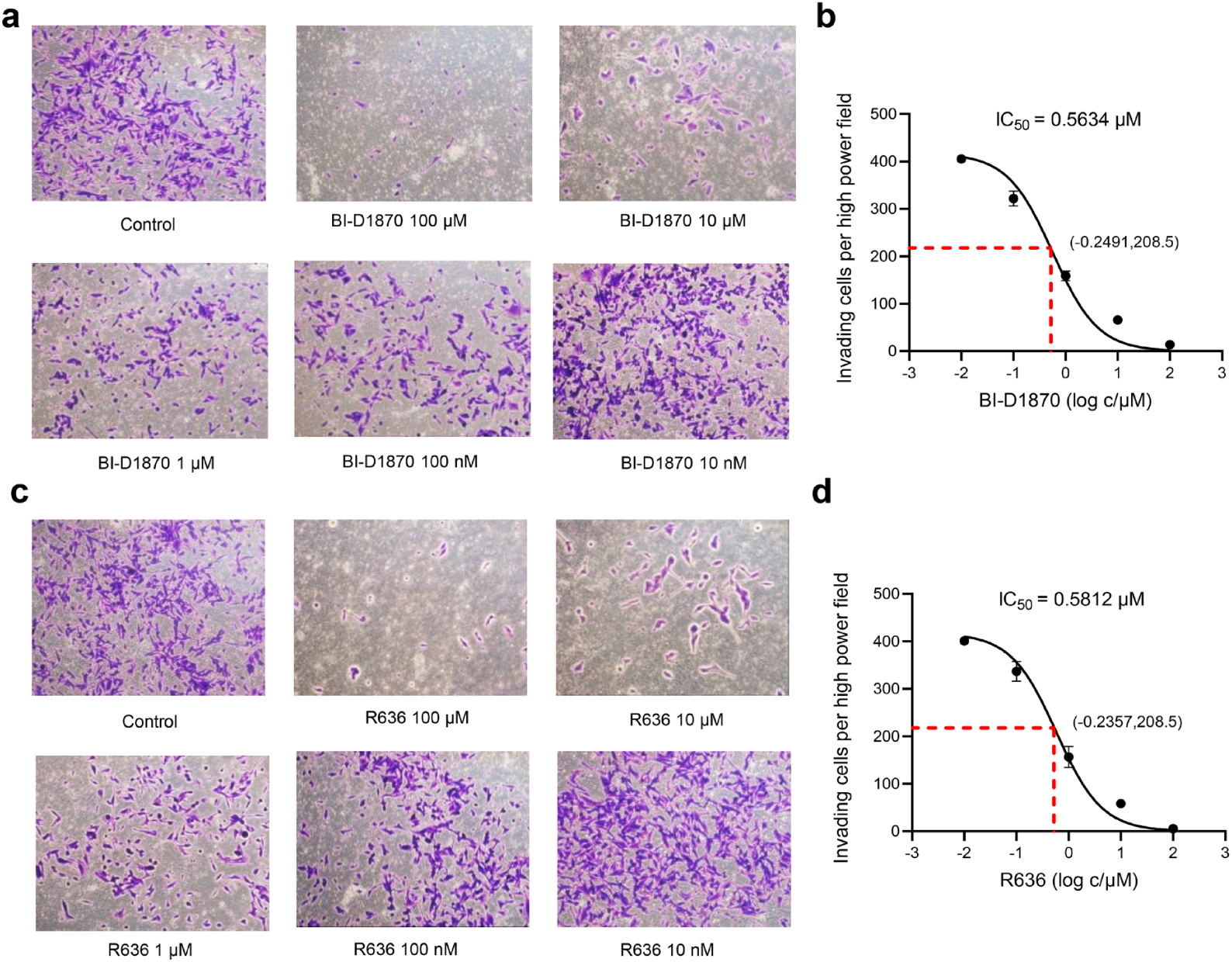
R636 inhibits TE10 cells’ invasion. Inhibitory activity of BI-D1870 (**a**, **b**) and R636 (**c**, **d**) against the invasion of TE10 cells in vitro by transwell analysis. TE10 cells were treated with various concentrations of BI-D1870 and R636 for 24 h.

**Extended Data Fig. 9.**
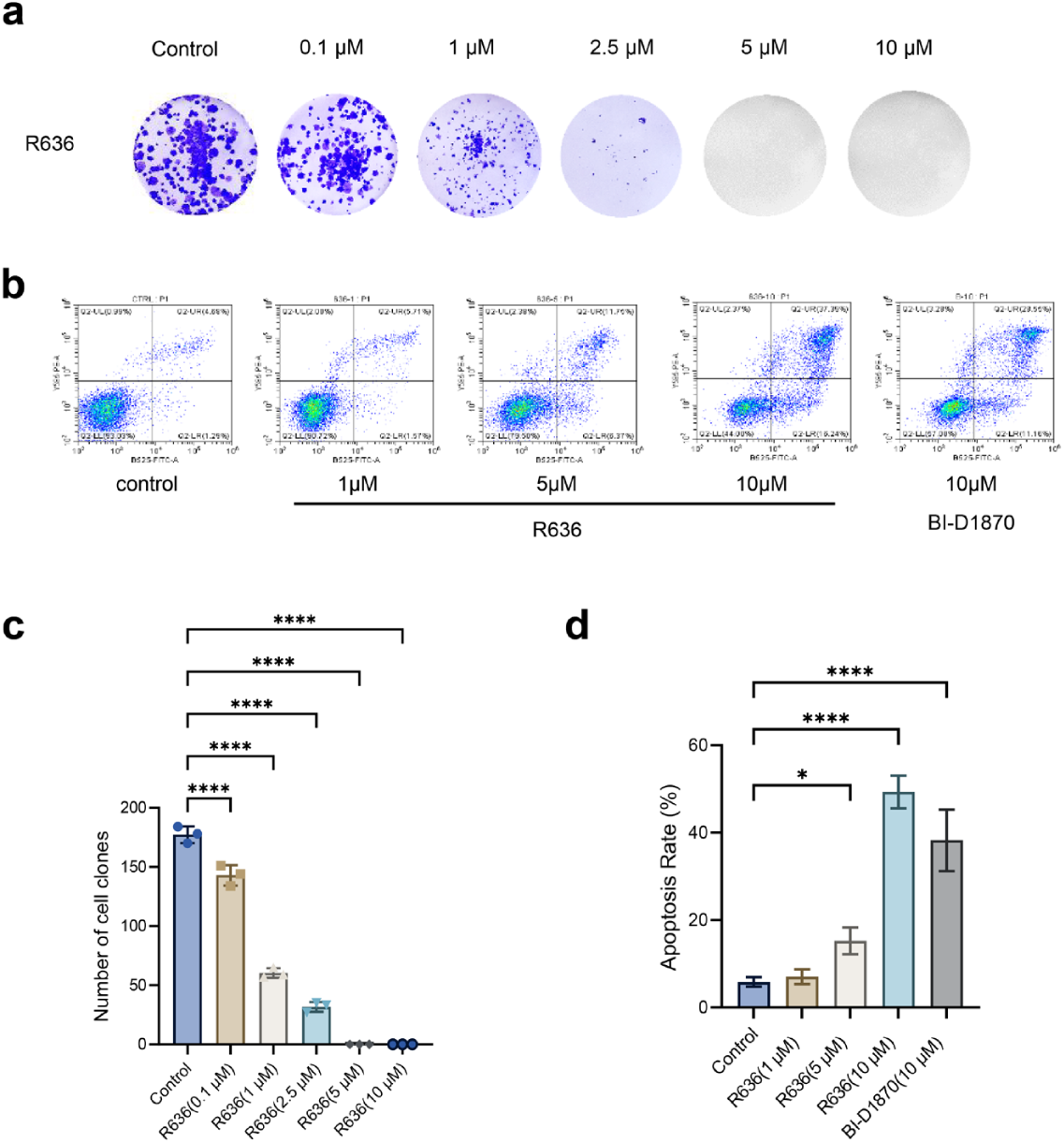
R636 suppresses colony formation and induces apoptosis in TE10 cells. **a**,**c**, R636 against the proliferation of TE10 cells by clone formation assay. **b**,**d**, R636 promotes apoptosis of TE10 cells.

**Extended Data Fig. 10.**
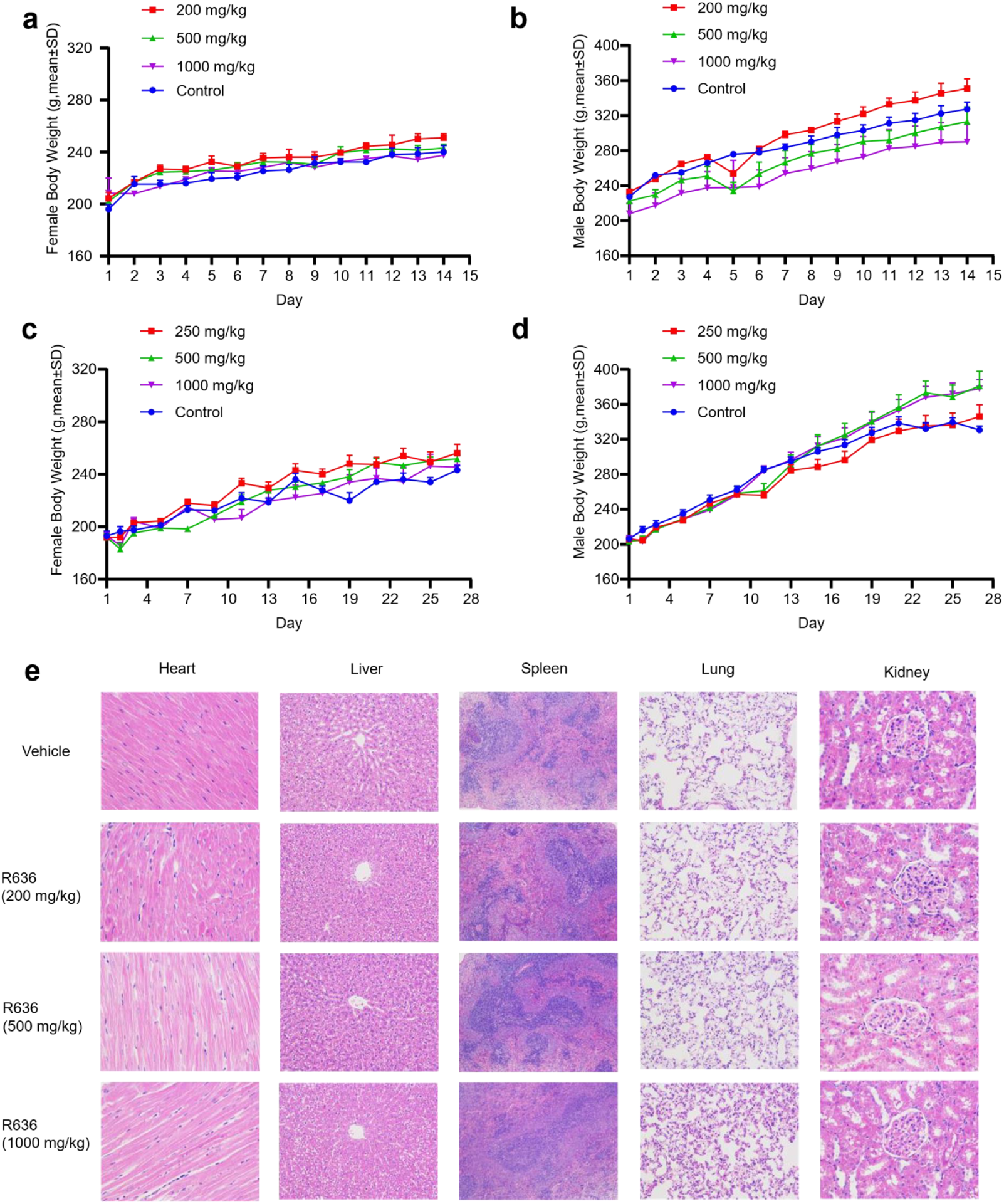
Acute and subacute toxicity evaluation of R636 in rats. a-b, The daily weight changes in each group (n = 4 Rats per group) in the acute toxicity experiment. c-d, The daily weight changes in each group (n = 8 Rats per group) in the subacute toxicity experiment. e, Representative image of HE analysis of main organs from the respective group, including heart, liver, spleen, lung and kindney in the subacute acute toxicity experiment.

**Extended Data Fig. 11.**
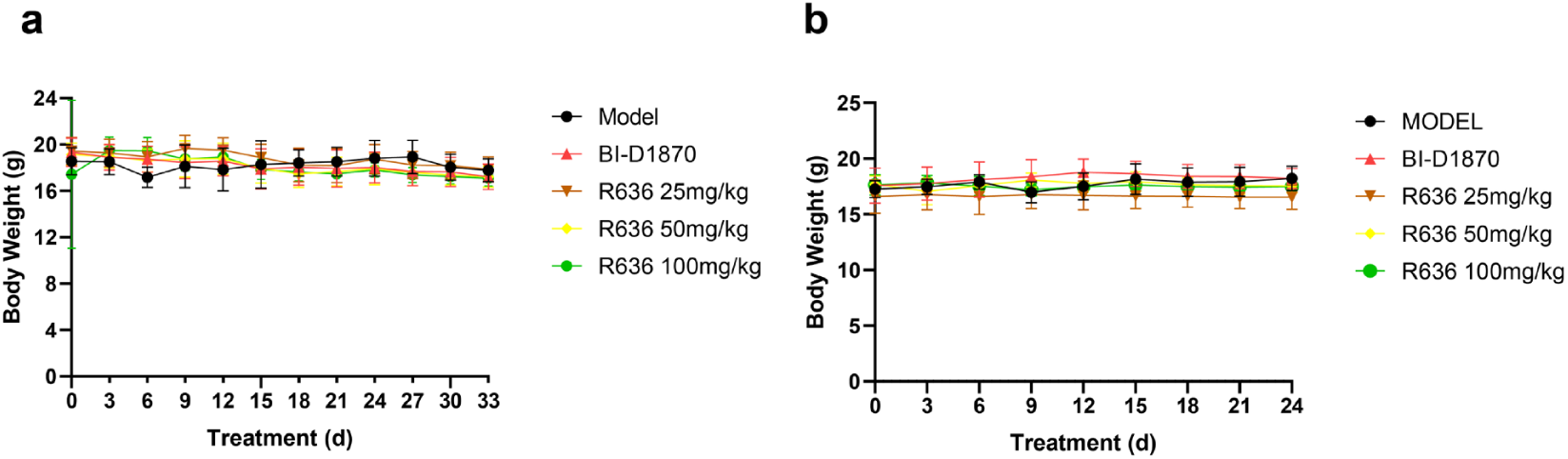
Body weight changes in mice during drug administration. Body weight changes in EG34 PDX model (**a**) and LEG110 PDX model (**b**).

## Supplementary Tables

**Supplementary Table 1.**
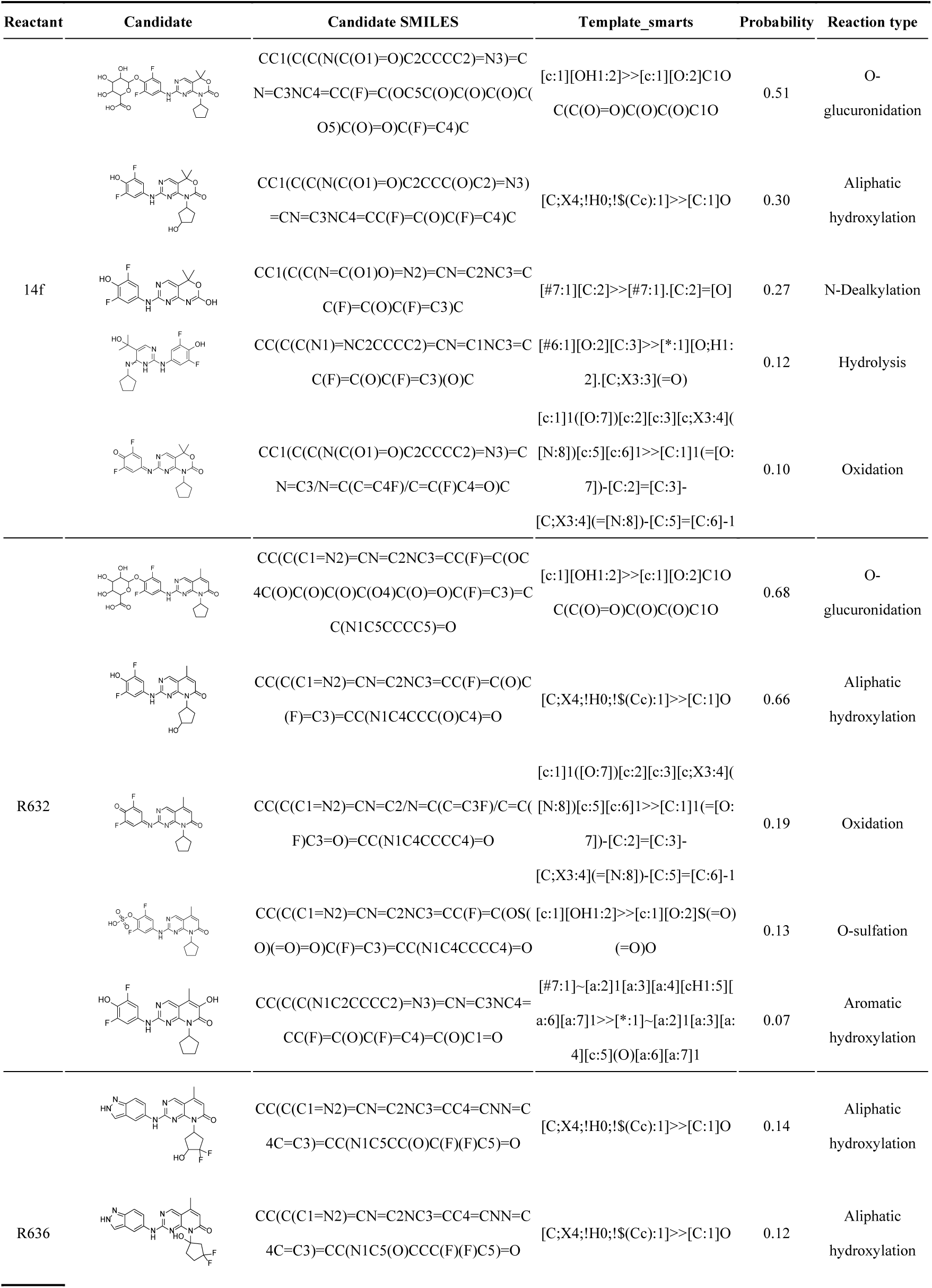
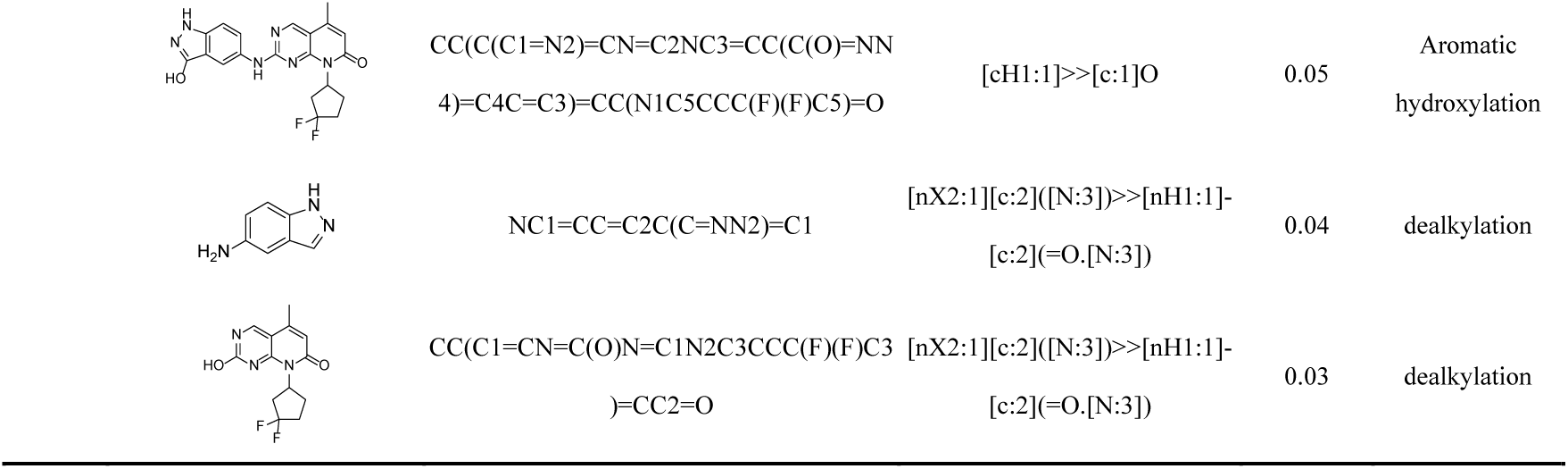
Metabolite prediction of 14f, R632, and R636.

**Supplementary Table 2.** Energy differences (Complex - Receptor - Ligand) for R632, R633 and R636.

| Differences (Complex - Receptor - Ligand) |  |  |  |  |  |
| --- | --- | --- | --- | --- | --- |
| R632 |  | R633 |  | R636 |  |
| Energy Component | Average | Energy Component | Average | Energy Component | Average |
| VDWAALS | -43.2762 | VDWAALS | -37.6089 | VDWAALS | -43.8805 |
| EEL | -30.9216 | EEL | -25.0717 | EEL | -23.451 |
| EGB | 37.4867 | EGB | 27.6572 | EGB | 31.2622 |
| ESURF | -5.4144 | ESURF | -4.4541 | ESURF | -5.3309 |
| DELTA G gas | -74.1978 | DELTA G gas | -62.6806 | DELTA G gas | -67.3315 |
| DELTA G solv | 32.0723 | DELTA G solv | 23.2031 | DELTA G solv | 25.9313 |
| DELTA TOTAL | -42.1255 | DELTA TOTAL | -39.4775 | DELTA TOTAL | -41.4002 |

<sup>a</sup>VDWAALS = van der Waals contribution. <sup>b</sup>EEL = electrostatic energy. <sup>c</sup>EGB = the electrostatic contribution to the solvation free energy calculated by GB. <sup>d</sup>ESURF = nonpolar contribution to the solvation free energy calculated by an empirical model. DELTA G<sup>e</sup> = final estimated binding free energy calculated from the terms above. (kcal/mol)

## Notes

### Competing Interest Statement

The authors have declared no competing interest.

